# *Pseudomonas aeruginosa* Elicits Sustained IL-1β Upregulation in Alveolar Macrophages from Lung Transplant Recipients

**DOI:** 10.1101/2022.04.26.489590

**Authors:** Noel Britton, Andres Villabona-Rueda, Samantha A. Whiteside, Joby Mathew, Matthew Kelley, Sean Agbor-Enoh, John McDyer, Jason D. Christie, Ronald G. Collman, Andrea Cox, Pali Shah, Franco D’Alessio

**Affiliations:** Department of Medicine, Johns Hopkins University, Baltimore, Maryland; Division of Pulmonary, Allergy, and Critical Care, Perelman School of Medicine, University of Pennsylvania, Philadelphia, Pennsylvania; Laboratory of Applied Precision Omics, National Heart, Lung, and Blood Institute, Bethesda, Maryland; Department of Medicine, University of Pittsburgh, Pittsburgh, Pennsylvania

## Abstract

**Background:** Isolation of *Pseudomonas aeruginosa* (*PsA*) is associated with increased BAL (bronchoalveolar lavage) inflammation and lung allograft injury in lung transplant recipients (LTR). However, the effect of *PsA* on macrophage responses in this population is incompletely understood. We examined human alveolar macrophage (AM) responses to *PsA and* Pseudomonas dominant microbiome in healthy lung transplant recipients (LTR).

**Methods:** We stimulated THP-1 derived macrophages (THP-1M) and human AM from LTR with different bacteria and LTR BAL derived microbiome characterized as Pseudomonas-dominant. Macrophage responses were assessed by high dimensional flow cytometry, including their intracellular production of cytokines (TNF-α, IL-6, IL-8, IL-1β, IL-10, IL-1RA, and TGF-β). Pharmacological inhibitors were utilized to evaluate the role of the inflammasome in *PsA*-macrophages interaction.

**Results:** We observed upregulation of pro-inflammatory cytokines (TNF-α, IL-6, IL-8, IL-1β) following stimulation by *PsA* compared to other bacteria (*Staphylococcus aureus, Prevotella melaninogenica, Streptococcus pneumoniae*) in both THP-1 derived and LTR AM, predominated by IL-1β. IL-1β production from THP-1 was sustained after *PsA stimulation* for up to 96 hours and 48 hours in LTR AM. Treatment with the inflammasome inhibitor BAY11-7082 abrogated macrophage IL-1β and IL-18 production after *PsA* exposure. BAL *Pseudom*onas-dominant microbiota elicited an increased IL-1β, similar to *PsA*, an effect abrogated by the addition of antibiotics.

**Conclusion:** *PsA* and *PsA*-dominant lung microbiota induce sustained IL-1β production in LTR AM. Pharmacological targeting of the inflammasome reduces *PsA*-macrophage-IL1β responses, underscoring their use in lung transplant recipients.

## Background

Lung transplantation is a life-extending treatment for individuals with end-stage lung diseases; however, long-term outcomes are limited by allograft injury.^1–5^ Infectious complications are among the strongest risk factors for acute and chronic lung allograft dysfunction (CLAD), the primary cause of mortality following lung transplantation.^6–9^

Lung allograft injury and CLAD are more frequent in lung transplant recipients (LTR) with clinical infection or colonization of the lung with specific pathogens, particularly de novo *Pseudomonas aeruginosa* ^10–12^ Emerging data suggest that increased bacterial burden, decreased bacterial diversity, and prominence of Proteobacteria, such as *PsA*, are associated with, graft failure, and CLAD through pathogen-driven inflammatory triggers and impaired innate responses impacting bacterial clearance. ^6– 8,11,13–15^ Regarding innate immune signaling, elevated IL-1β has been associated with a decline in lung function, injury in murine models of infection, and the development of CLAD; ^16–21^ but, few studies have examined the role of pathogen-driven inflammatory triggers on human innate immune responses in lung transplantation.^6–8,11,13–15^ Recent work has shown that Proteobacteria can enhance inflammatory M1 macrophage activation with excess inflammation or tissue injury and that resolution of Pseudomonas-lung injury in other lung disease models is mediated by a shift of macrophage responses.^22–24^ However, the role of *PsA* in the lung microbiome and its activation of alveolar macrophage (AM) innate immune signaling following lung transplantation is incompletely understood.

While CLAD is defined as a generally irreversible clinical phenotype of lung injury, the antecedent risk factors may occur months or years prior to diagnosis, supporting a hypothesis that early inflammatory mechanisms could represent important triggers for CLAD. ^25^ Given the association of *PsA*, microbiome shifts, and IL1-β with CLAD, we sought to examine the innate immune response of macrophages following exposure to *PsA* or *PsA* dominant microbiota. As such, we hypothesized that the lung microbiome, particularly those dominated by *PsA*, following transplantation mediates sustained AM proinflammatory responses. We determined that *PsA* predominantly upregulates IL-1β production in AM compared to other pathogenic and commensal airway bacteria. We show that this effect is sustained and that macrophage IL-1β responses to *PsA* can be abrogated using BAY11-7082, an inflammasome inhibitor. Our data from both a preclinical model system and *ex vivo* AM from stable LTR suggest that inflammasome-mediated IL-1β responses may represent an important mechanistic pathway in PsA-associated inflammation following lung transplantation.

## Methods

### Study Approval

Consent was obtained for all participants in this prospective observational study. All human work was approved by the IRB at the Johns Hopkins Hospital (IRB00138643, IRB00236519) or the University of Pennsylvania (#812748).

### Study Population

Eligible participants were undergoing surveillance bronchoscopy between 12-24 months post-transplant without suspicion of active rejection or infection (Supplementary Table 1). Participants were on the institution’s baseline immunosuppression and infectious disease prophylaxis (Supplementary Methods).

### BAL Cell Collection

BAL was performed per ISHLT guidelines and processed as previously described.^26^ We centrifuged BAL fluid at 300 g for 10 minutes to pellet cells. Cells were maintained in supplemented tissue culture medium (Roswell Park Memorial Institute (RPMI) 1640 media with 10% FBS, 2mM L-Glutamine, and 50 μM β-mercaptoethanol, 100 U/mL penicillin/streptomycin) and plated 0.5-1×10^6^ cells per sample at 37 °C, 5% CO_2_ for stimulation. For bacterial stimulation, cells were stimulated with individual bacterial species at a multiplicity of infection of 1. Cells were stimulated for 24 and 48 hours.

### THP-1 Derived Macrophages

The human monocyte-like cell line THP-11 (TIB-202) was obtained from the American Type Culture Collection (ATCC). Cells were grown in supplemented tissue culture medium in the range of 2-9×10^5^ cells/ml at 37 °C, 5% CO_2_. We incubated THP-1 cells in non-treated 24 well cell culture plates with phorbol 12-myristate 13-acetate (PMA) 100 ng/mL for 24 hours to differentiate cells into macrophages. The cells were washed with phosphate-buffered saline (PBS), cultured in supplemented tissue culture medium without antibiotics, and plated 1×10^6^ cells per well for stimulation with culture media (control), lipopolysaccharides from Escherichia coli O55:B5 (Sigma-Aldrich #L2880) at a concentration of 100 ng/µL.and bacterial species at a multiplicity of infection of 1, for 24, 48, 72, and 96 hours.

### Bacterial Preparation

Experimental stock cultures of *Pseudomonas aeruginosa, Prevotella melaninogenica, Staphylococcus aureus*, and *Streptococcus pneumoniae* were prepared as described in the supplement. For stimulation, bacteria were thawed by gentle agitation in a 37°C water bath, centrifuged at 130g for 10 minutes at 20°C to pellet cells, and then resuspended in tissue culture medium. For heat-inactivated stimulations, thawed bacteria were inactivated at 60°C for 60 minutes.

### Microbiome Stimulation

BAL was collected using ISHLT guidelines with methods specific for microbiome analysis from LTR (Supplementary Table 2).^26–28^ DNA extraction, 16S rRNA gene PCR amplification using V1V2 primers (27F and 338R), and Illumina sequencing on the MiSeq platform was performed as previously described.^29,30^ Processing of sequencing data is detailed in the Supplement. Bacterial load was assessed by qPCR targeting the 16S rRNA gene.^26^ The volume of BAL used to stimulate cells was standardized to a bacterial burden equivalent to a concentration of 0.05 pmol/µL.16s rRNA. THP-1M were stimulated with *PsA*-dominant BAL for 24 hours.

### Inflammasome Inhibitors

THP-1M were stimulated with *PsA* for 24 hours. Inflammasome inhibitors BAY11-7082, glybenclamide, parthenolide, and MCC950 (Invivogen) were administered as preventive (1 hour before *PsA* stimulation) or rescue (6 hours after *PsA* stimulation) at a concentration of 10 μM.

### Flow Cytometry

Following stimulation, cells were stained for cell surface markers and intracellular cytokines as described in the Supplement. Cells were then analyzed on a FACSAria (BD Biosciences) and FACSDiva for data acquisition (BD Biosciences). Data were analyzed using FlowJo (TreeStar, Inc.) for gating analyses. THP-1M were first gated for single cells, and then dead cells were gated out. To interrogate LTR AM specifically from BAL cells, single cells were gated out for dead cells and then gated for CD206+CD11b+CD64+ macrophages.

### ELISA

Macrophage IL-18 was measured with the human IL-18 ELISA Kit electrochemiluminescence assay (MBL International) per manufacturers’ instructions using cell culture supernatant at a 1:5 dilution. Data were acquired on a SpectraMax M2.

### Statistics

*In vitro* experiments were performed in triplicate and repeated at least three times. Mean fluorescence intensity (MFI) was calculated as the mean of the positive population fluorescence. Differences between groups were compared using a two-tailed nonparametric Wilcoxon test. Multiple group comparisons were performed using a Kruskal Wallis test with Tukey’s multiple comparisons test. A *P* value less than 0.05 was accepted as significant. Statistics were performed using R (RStudio 2021.09.2+382 “Ghost Orchid”) and Prism (Version 9.3.1, GraphPad).

## Results

### Macrophage pro-inflammatory responses to different bacterial taxa

To identify the differential effects of unique bacterial species on macrophage immunophenotype, we stimulated THP-1M for 24 hours with bacteria representing infectious pathogens observed in the lung (e.g., *PsA, Staphylococcus aureus)* and with bacteria representing upper airway commensals (e.g., *Prevotella melaninogenica, Streptococcus pneumoniae*). Bacteria were heat-inactivated to avoid differential bacterial replication.

We observed diverse macrophage immune sensing and responses to different bacteria, most notably, preferential expression of IL-1β, an important inflammasome mediator, in response to *PsA*. Expression of IL-1β (ΔMFI 1.75, p<0.001) and IL-8 (ΔMFI 1.30, p=0.02) were elevated in THP-1M stimulated with *PsA* compared to unstimulated controls. Compared to controls, we did not observe significant differences in proinflammatory TNF-*α*, anti-inflammatory IL-RA or IL-10 expression in macrophages stimulated with bacteria but did observe anti-inflammatory TGF-β elevated in *PsA-*stimulated THP-1M (ΔMFI 0.92, p<0.004) (**Figure 1A)**.

**Figure 1:**
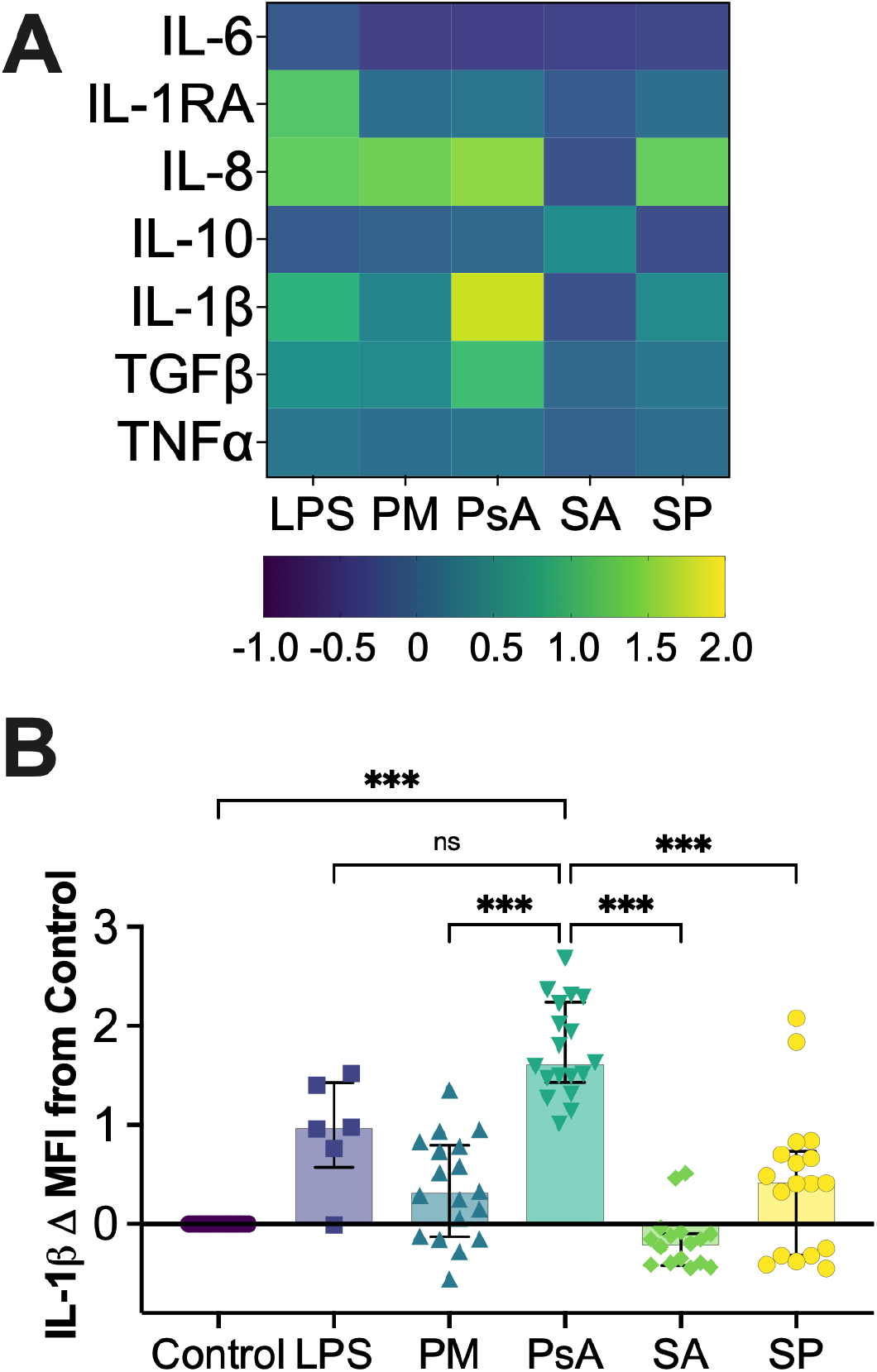
THP-1MΦ; responses to different bacteria at 24 hours. THP-1MΦ (n=18) were stimulated with heat inactivated bacteria at a multiplicity of infection of 1:1 for 24hours. Cells were incubated with Golgi inhibitors the last 4 hours, stained for high dimensional flow cytometry. Each marker is expressed as mean fluorescence intensity (MFI) fold change compared to expression in macrophages at baseline conditions (control). **A)** Heatmap for THP-1MΦ cytokine expression after bacterial stimulation for 24 hours. **B)** THP-1MΦ express elevated IL-1β in response to *PsA* compared to other bacteria. **ns** = not significant; **\*** p ≤0.05; **\*\*** p ≤0.002; **\*\*\*** p ≤ 0.001; **LPS**= lipopolysaccharide, **PM**= P. melaninogenica, **PsA**=P. aeruginosa, **SA**= S. aureus, **SP**= S. pneumoniae

While the overall pro-inflammatory response of THP-1M was elevated following 24 hours of *PsA* stimulation, IL-1β was most notably upregulated following *PsA* stimulation compared to unstimulated controls and to macrophages stimulated with other bacteria (p<0.001) (**Figure 1B)**. THP-1M had sustained IL-1β after stimulation with PsA compared to unstimulated macrophage (controls) at 48 (ΔMFI 1.41, p<0.001), 72 (ΔMFI 1.38, p<0.001), and 96 (ΔMFI 1.21, p<0.001) hours (**Figure 2)**. IL-1β production by THP-1M after *PsA* remained elevated after 96 hours compared to control macrophages in response to *S. aureus, P. melaninogenica*, and *S. pneumoniae* (**Supplement Figure 1)**.

**Figure 2:**
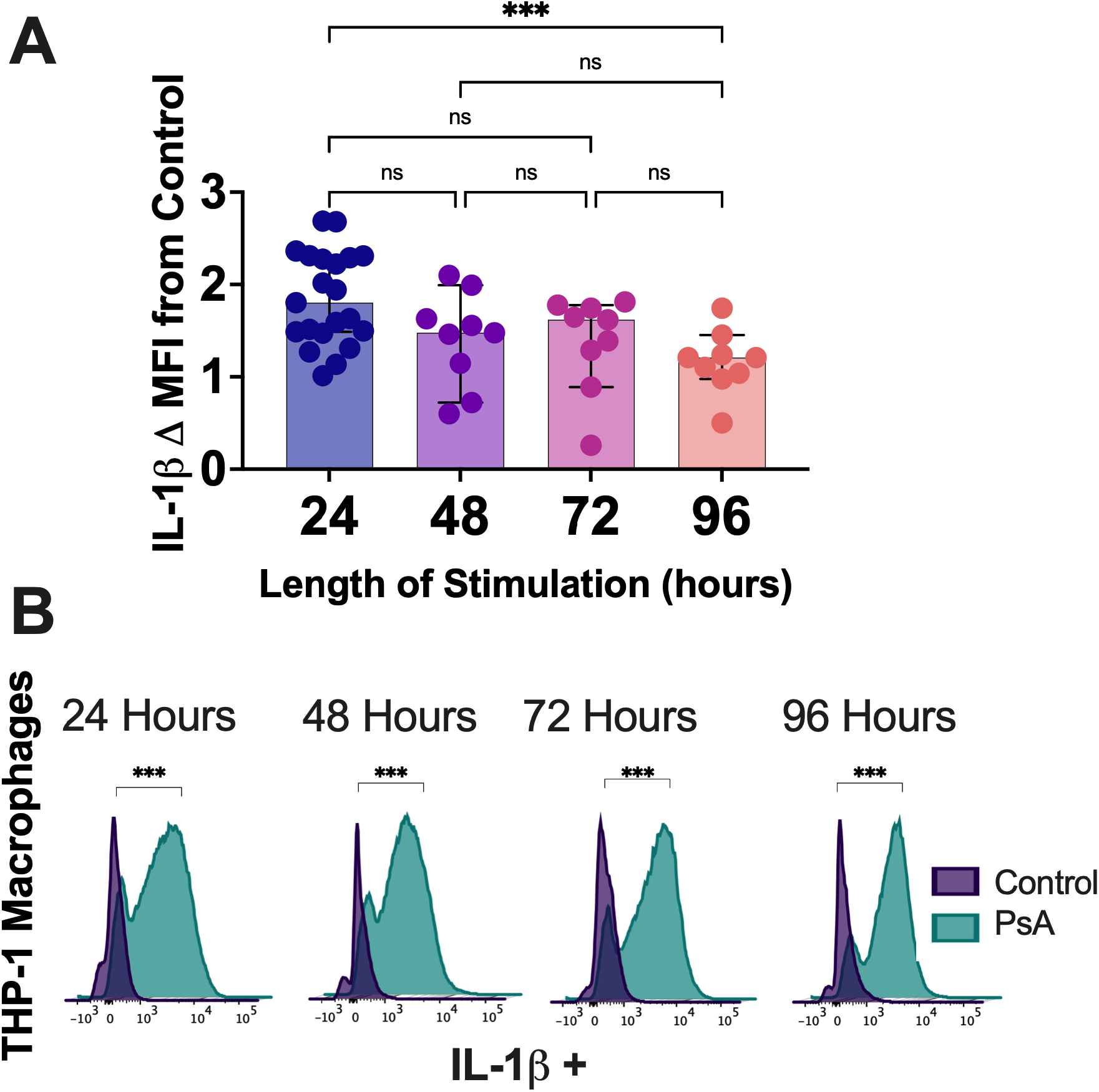
THP-1MΦ stimulated with PsA had sustained elevation of IL-1β species over time. THP-1MΦ were stimulated with *PsA* at a multiplicity of infection of 1:1 for 24(n=21), 48 (n=9), 72(n=9) and 96(n=9) hours. Cells were incubated with Golgi inhibitors the last 4 hours and stained for high dimensional flow cytometry. **A)** IL-1β kinetics after challenge with *PsA* for up to 96 hours are displayed as mean fluorescence intensity (MFI) fold change (Δ) compared to expression in macrophages at baseline conditions (control). **B)** Representative histograms demonstrating IL-1β expression after *PsA* stimulation compared to unstimulated controls for 24, 48, 72 and 96 hours. **ns** = not significant; **\*** p ≤0.05; **\*\*** p ≤0.002; **\*\*\*** p ≤ 0.001

### Alveolar macrophage response to bacteria is comparable to THP-1 macrophages

We used human LTR AM to determine that the responses seen in our cell line translated into primary human cells. *Ex vivo* BAL-derived cells from LTR were stimulated with diverse bacteria as with THP-1 macrophages. Baseline clinical characteristics for the human subjects enrolled in this observational study are shown in (**Supplement Table 1**). LTR AM (CD64+CD206+CD11b+) responses demonstrated an overall upregulation of pro-inflammatory cytokines following 24 hours of stimulation by *PsA* compared to stimulation with *S. aureus, P. melaninogenica, S. pneumoniae (***Figure 3A)**, similar to our earlier observations in THP-1 cell lines. Notably, there was increased production of IL-1β by AM after exposure to *PsA* compared to macrophage control (ΔMFI 2.24, p<0.001), P. *melaninogenica*, and *S. pneumoniae*, but not *S. aureus* **(Figure 3B**). Parallel to responses in THP1-M, we observed sustained IL-1β production in LTR AM following stimulation by *PsA* at 48 hours (**Figure 4**) compared to controls (ΔMFI 2.52, p <0.001), and LTR AM stimulated with *PsA* demonstrated higher IL-1β expression than those stimulated with other bacteria at 48 hours (p<0.001) (**Supplement Figure 2**).

**Figure 3:**
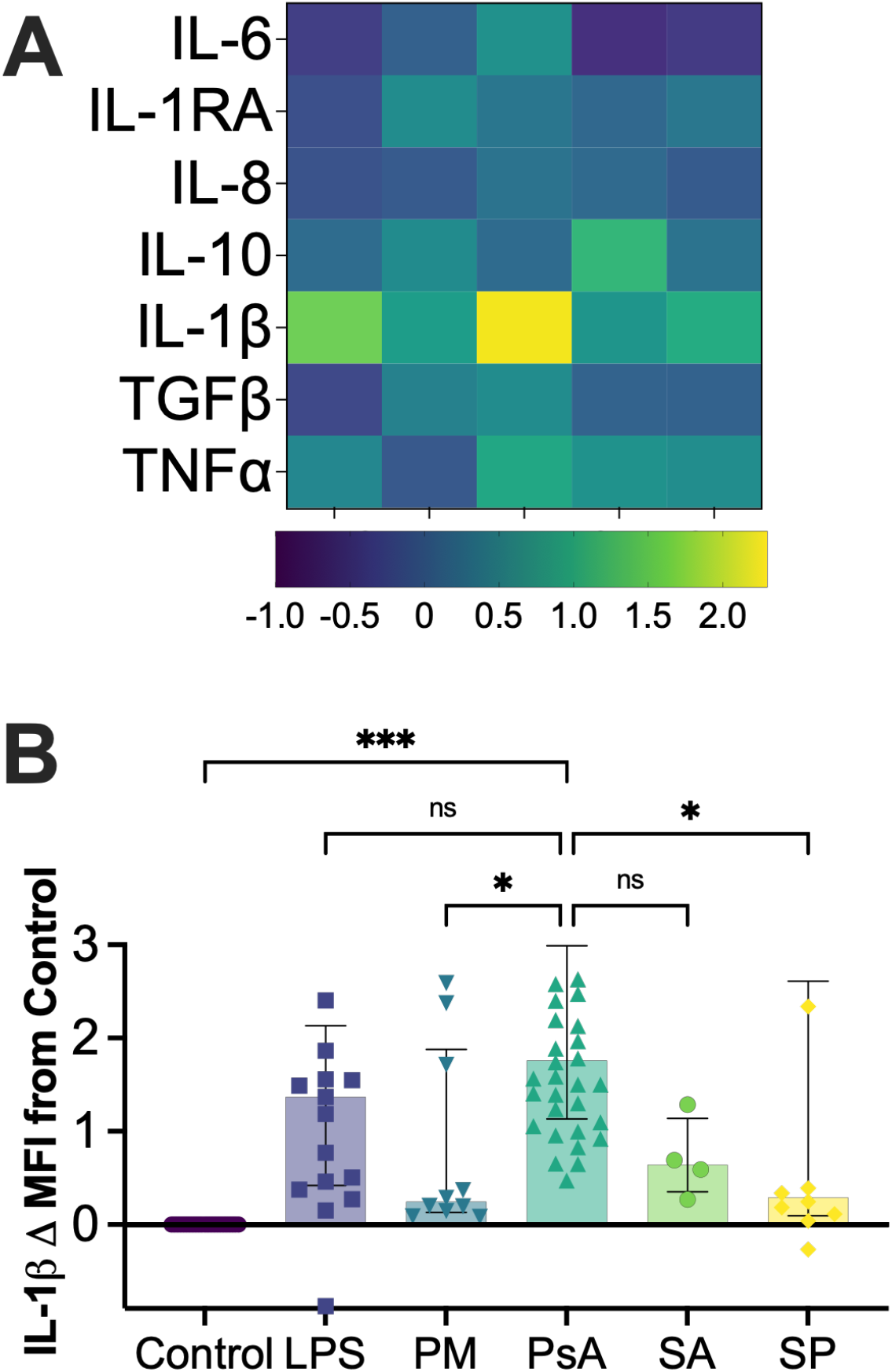
AMΦ responses to different bacteria at 24 hours. *AMΦ* were stimulated with heat inactivated bacteria at a multiplicity of infection of 1:1 for 24hours. Cells were incubated with Golgi inhibitors the last 4 hours, stained for high dimensional flow cytometry. Each marker is expressed as mean fluorescence intensity (MFI) fold change compared to expression in macrophages at baseline conditions (control). **A)** Heatmap for *AMΦ* cytokine expression after bacterial stimulation for 24 hours. **B)** *AMΦ* express elevated IL-1β in response to *PsA* compared to other bacteria. **LPS** (n=17), **PM** (n=10), **PsA** (n=20), **SA** (n=5), **SP** (n=10); **ns** = not significant; **\*** p ≤0.05; **\*\*** p ≤0.002; **\*\*\*** p ≤ 0.001; **LPS**= lipopolysaccharide, **PM**= P. melaninogenica, **PsA**=P. aeruginosa, **SA**= S. aureus, **SP**= S. pneumoniae

**Figure 4:**
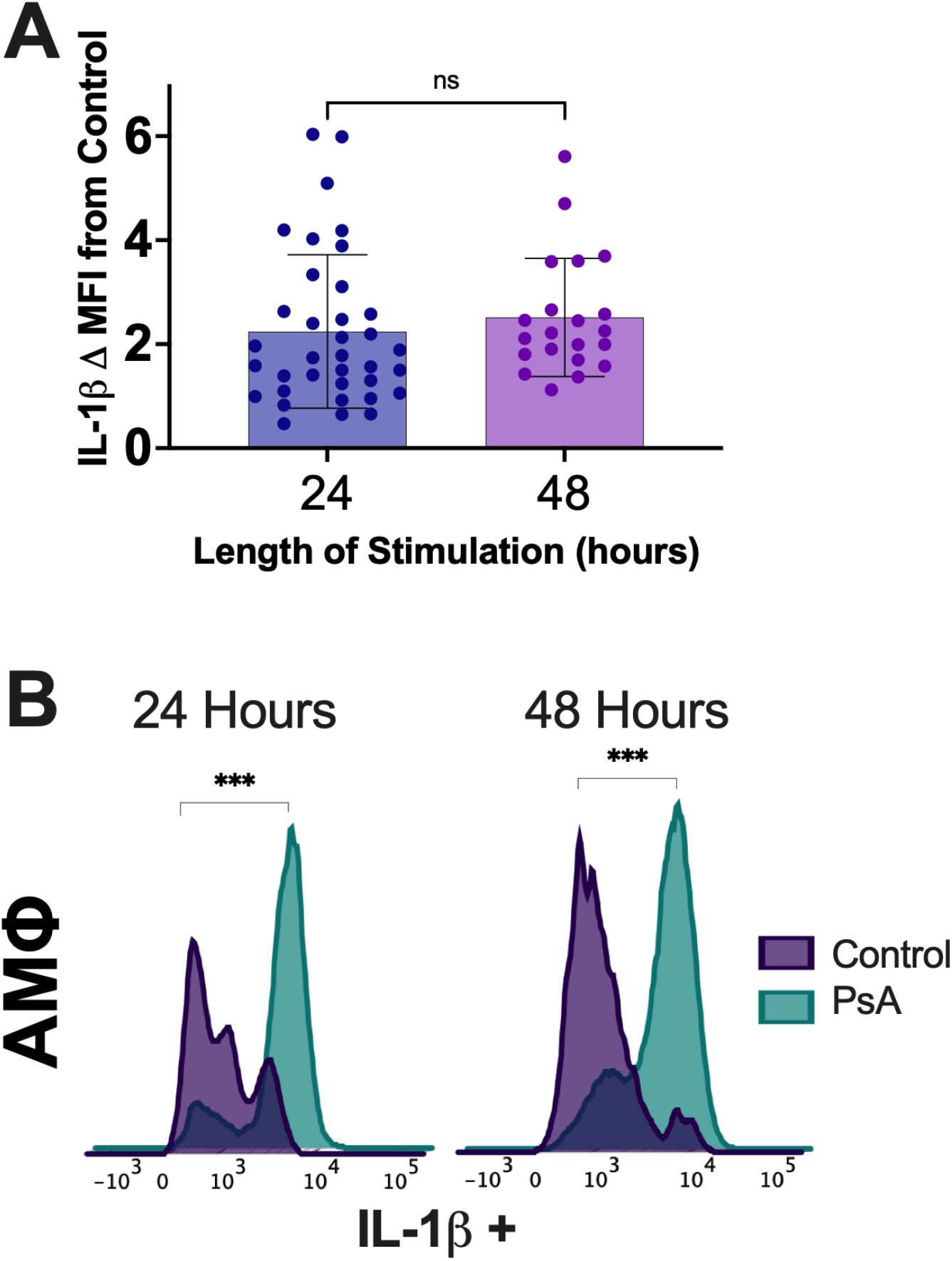
AMΦ stimulated with PsA had sustained elevation of IL-1β species over time. *AMΦ* were stimulated with *PsA* at a multiplicity of infection of 1:1 for 24(n=36) and 48 (n=21) hours. Cells were incubated with Golgi inhibitors the last 4 hours and stained for high dimensional flow cytometry. **A)** IL-1β kinetics after challenge with *PsA* for up to 48 hours are displayed as mean fluorescence intensity (MFI) fold change (Δ.) compared to expression in macrophages at baseline conditions (control). **B)** Representative histograms demonstrating IL-1β expression after *PsA* stimulation compared to unstimulated controls for 24 and 48 hours. **ns** = not significant; **\*** p ≤0.05; **\*\*** p ≤0.002; **\*\*\*** p ≤ 0.001

### Inflammasome inhibition prevents and rescues *PsA*-induced IL-1β production in macrophages

Given that secretion of IL1β is regulated by activation of the inflammasome, we sought to investigate if we could modulate macrophage inflammasome responses to *PsA*. We first pre-treated THP-1M with inflammasome inhibitors designed to inhibit NLRP3 and NLPC4 inflammasome pathways 1 hour prior to stimulation with PsA. 24 hours after stimulation, we observed a striking magnitude of reduction in IL-1β in THP-1M pre-treated with BAY11-7082, a multi-target NF-κB pathway inhibitor that also inhibits NLRP3 and may partially inhibit the NLRC4 inflammasome, at 24 hours after *PsA* (**Figure 5A**), to a level below what was constitutively expressed by control unstimulated macrophages. Treatment with parthenolide, a multi-target inhibitor of NF-κB, NLRC4, and NLRP3, and glybenclamide, an NLRP3 specific inflammasome inhibitor, reduced IL-1β expression to levels similar to those observed in unstimulated macrophage control**s**. To evaluate if IL-1β production could be reduced after exposure to *PsA*, we rescue-treated AM inflammasome inhibitors 6 hours after stimulation and measured IL-1β levels 24 hours post stimulation. Treatment with rescue BAY11-7082, parthenolide, and glybenclamide, reduced IL-1β expression to levels like those observed in preventive dosing. (**Figure 5B**).

**Figure 5:**
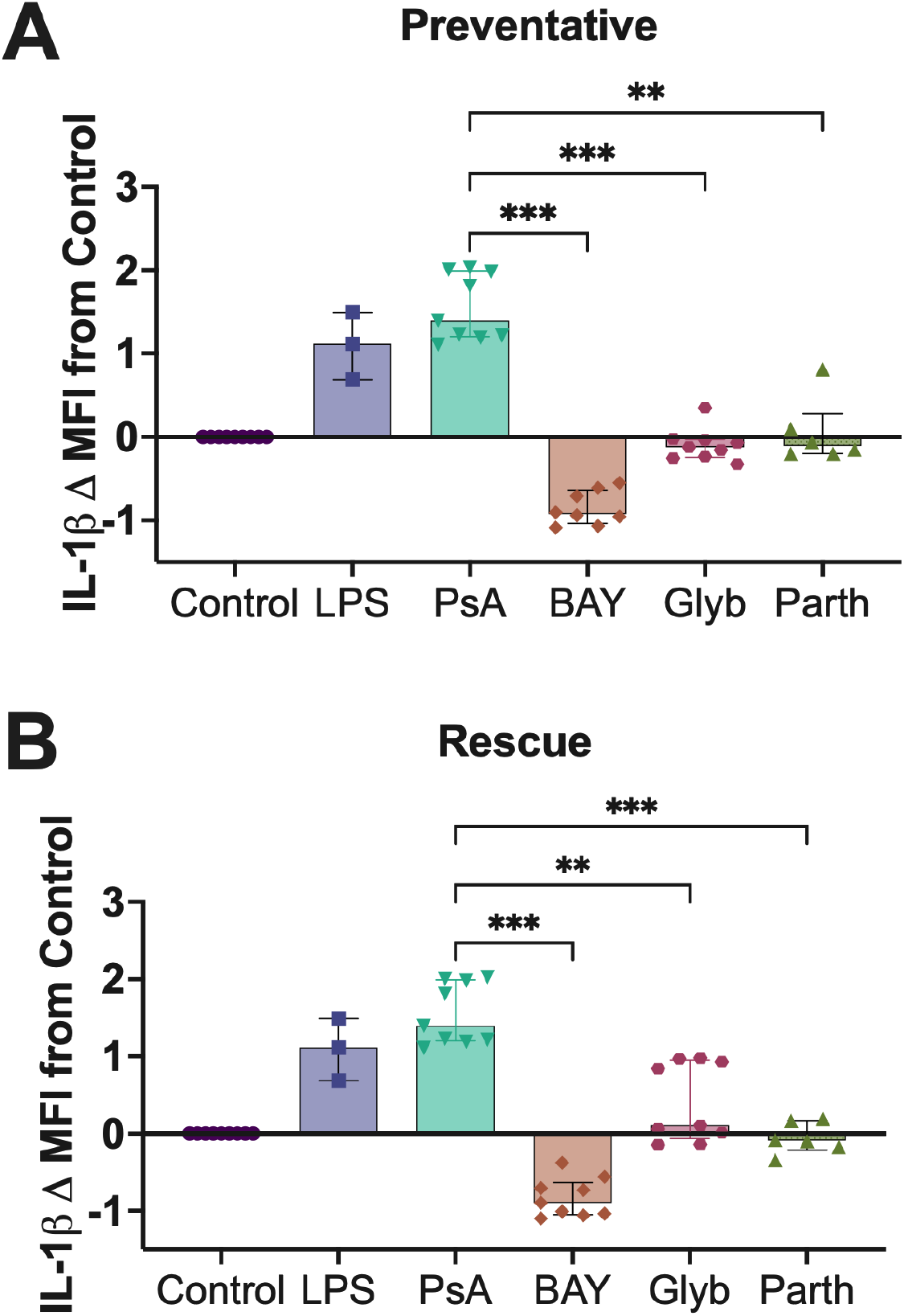
Inflammasome pharmacological inhibition in macrophage-PsA interactions. THP-1MΦ were stimulated with *PsA* treated with inflammasome inhibitors in a preventative or rescue dosing schedule for 24 hours. THP-1MΦ were treated with Golgi inhibitors the last 4 hours of the 24 hour stimulation and were stained for high dimensional flow cytometry. THP-1MΦ were then interrogated for their relative expression of IL-1β expression. IL-1β is expressed as mean fluorescence intensity (MFI) fold change compared to expression in macrophages at baseline conditions (control). **A)** THP-1MΦ treated with a preventative dosing schedule were first treated with inflammasome inhibitors and then stimulated with *PsA* after 6 hours. **B)** THP-1MΦ treated with a rescue dosing schedule were first stimulated with *PsA* and then treated with inflammasome inhibitors. **ns** = not significant; **\*** p ≤0.05; **\*\*** p ≤0.002; **\*\*\*** p ≤ 0.001; **LPS**= lipopolysaccharide, **PsA**= P. aeruginosa, **BAY**= BAY11-7082, **Glyb**= glybenclamide **Parth**=parthenolide

### *PsA* dominant lung microbiome communities from LTR elicit robust IL-1β upregulation in macrophages

To determine if lung microbiota samples from LTR that were *Pseudomonas* dominated would comparably stimulate macrophages, we exposed THP-1M for 24 hours with BAL-derived microbiota from three LTR that had ≥90% relative abundance of *PsA*. Following lung microbiota stimulation, we observed production of IL-1β comparable to macrophages challenged with single species *PsA* (p=0.17) (**Figure 6**). The addition of antibiotics (Penicillin/Streptomycin) (p<0.001), but not antifungals (p=0.51), partially abrogated IL-1β responses suggesting that these responses are driven mainly by bacterial taxa.

**Figure 6:**
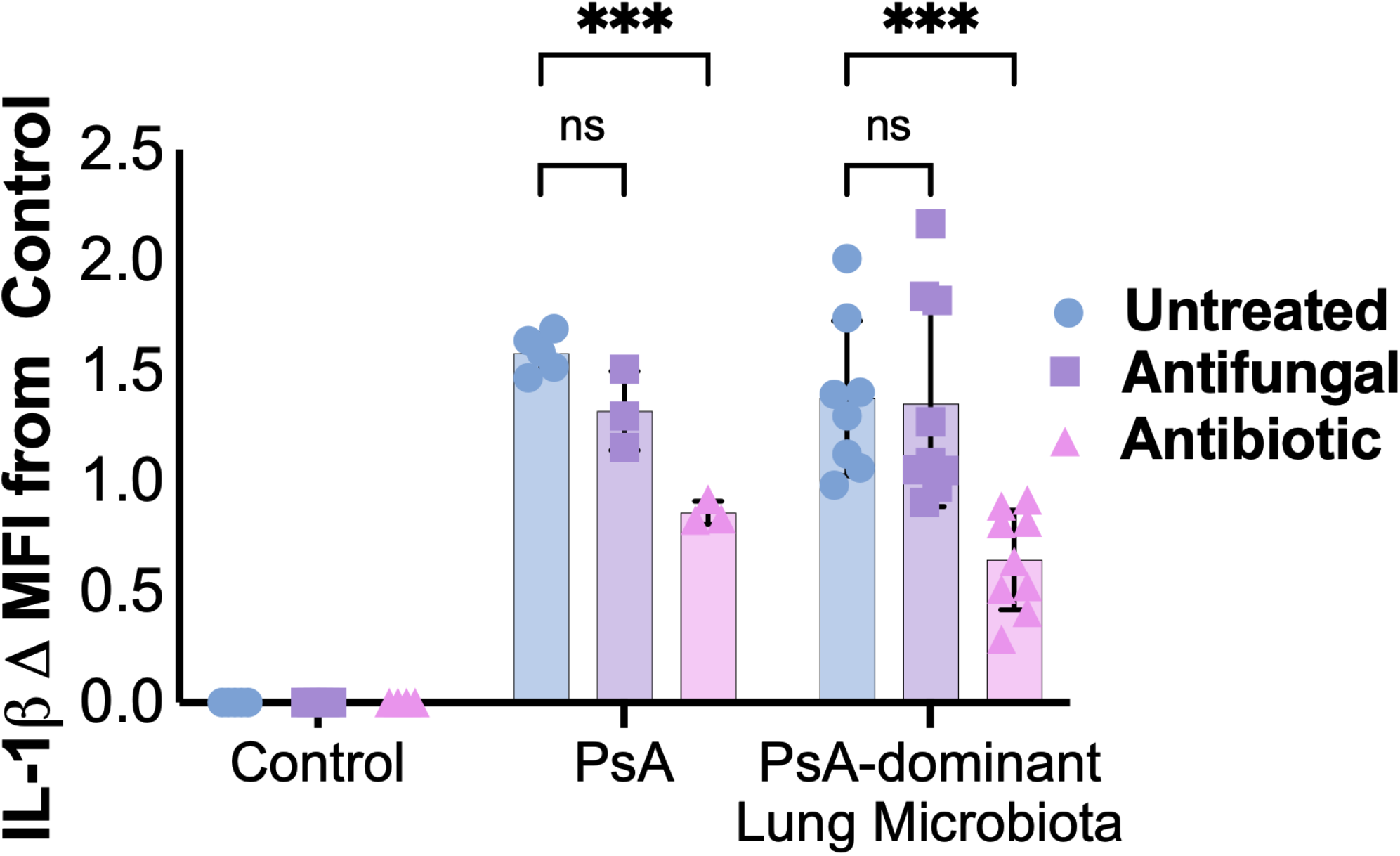
THP-1MΦ IL-1β upregulation after Pseudomonas-dominant LTR BAL microbiome stimulation for 24 hours. LTR BAL microbiota exposure was standardized using 16S qPCR measurement of bacterial burden. THP-1MΦ (n=9) were stimulated with eukaryotic cell-strained LTR BAL microbiota with a relative abundance greater than 90% *Pseudomonas* calculated by 16S sequencing for 24 hours. THP-1MΦ were treated with Golgi inhibitors the last 4 hours of the 24 hour stimulation and were stained for high dimensional flow cytometry. IL-1β is expressed as fold change in MFI compared to in macrophages at baseline conditions (control). THP-1 macrophages stimulated with *Pseudomonas-*dominant LTR BAL microbiota were treated with a broad-spectrum antifungal. THP-1 macrophages stimulated with *Pseudomonas-*dominant LTR BAL microbiota were also treated with both an antifungal and an antibiotic. Differences in IL-1β expression between untreated, antifungal-treated and antifungal antibiotic treated THP-1MΦ stimulated with LTR BAL microbiota were examined. **ns** = not significant; **\*** p ≤0.05; **\*\*** p ≤0.002; **\*\*\*** p ≤ 0.001

## Discussion

Our studies found that *PsA* exposure led to a sustained upregulation of pro-inflammatory cytokine production by AM, predominantly IL-1β, in both cell line and human lung transplant AM. We saw a similar upregulation of IL-1β in macrophages stimulated with *Pseudomonas-dominant* BAL-derived lung microbiota and demonstrated that treatment with inflammasome inhibiting drugs ameliorates this upregulation. Taken together, our findings provide evidence for inflammasome activation as a potential mechanistic link between *Pseudomonas*-driven microbial dysbiosis and sustained innate immune activation by AM from otherwise healthy LTR. Given prior reports of IL-1β upregulation in lung injury post-transplant, our data provide an important mechanism to further investigate the downstream development of CLAD.

Compared to bacteria representative of other pathogens and upper respiratory commensals, *PsA* was responsible for a sustained upregulation of pro-inflammatory cytokines by THP-1 and human AM, specifically IL-1β. Clinical studies have demonstrated the longitudinal persistence of *PsA in* LTR, and thus our findings suggest that this could lead to an aberrant and persistent upregulation of IL-1β. In parallel, it has been shown that IL-1β expression is elevated in LTR with *PsA* infection and colonization and that BAL IL-1β levels are increased prior to the development of and in LTR with CLAD. In prior studies, stimulations of cell line derived macrophages were performed using single bacteria stimulations to validate the associations identified between *Pseudomonas* and lung injury but have been limited in the direct assessment of human AM from LTR.^13,31^ Our results support other findings examining gene expression arrays of BALF from LTR, which suggest catabolic parenchymal remodeling and distinct inflammatory profiles are associated with lung microbiota communities dominated by inflammatory-associated bacteria, including *Pseudomonas*. ^31^ Our novel studies support a critical mechanistic role of macrophage-associated inflammation in the lung allograft.^32,33^

We saw a similar upregulation of IL-1β in macrophages stimulated with *Pseudomonas*-dominant BAL-derived lung microbiota. To the best of our knowledge, these are the first set of studies using BAL microbiome as stimuli to evaluate macrophage sensing and innate immune responses. Studies involving germ-free mice have previously shown a critical role of the microbiome in regulating the inflammasome and its activation, including downstream IL-1β and IL-18 responses.^34^ BAL samples from LTR with elevated concentrations of IL-1β have been associated with lower microbial diversity and were compositionally different than samples with lower concentrations of IL-1β but without identifying targeted cellular pathways.^19^ While the observation of macrophage activation by Pseudomonas-dominant LTR BAL communities *ex vivo* is concordant with the activation triggered by purified Pseudomonas bacteria, further studies will be helpful to delineate other potential contributors within the complex human LTR lung microbiome, including fungal and viral components.

IL-1β mediates a wide range of immune and inflammatory responses and is regulated through inflammasome activation by microbial and danger associated molecular patterns. ^35^. *Pseudomonas* infection of the lung has been associated with increased expression of inflammasome sensors NLRP3 and NLRC4 ^36 17^. Thus, inflammasome activation by *PsA* through the NLRC4 and NLRP3 sensors may represent a mechanistic link between *PsA* and immune responses in the allograft. We examined if pharmacological blockade of the inflammasome pathway could represent a therapeutic target for the sustained IL-1β production from macrophage-*Pseudomonas* interaction. We observed that *PsA* stimulated IL-1β and IL-18 production was diminished in macrophages treated with pan-inflammasome inhibitors compared as well as agents that preferentially targeted NLRP3, supporting the potential for using targeted inflammasome inhibitors to ameliorate chronic inflammation caused by sustained IL-1β production.

Persistent inflammasome activation plays a major role in lung inflammation and has been implicated in several chronic pulmonary diseases associated with chronic inflammation and fibrosis and autoimmune conditions.^37–40^ The sustained and aberrant activation of the inflammasome we describe may modulate microbiota-induced inflammation, and should future studies confirm causality with lung allograft injury; this pathway could be a potential therapeutic avenue for lung transplantation, particularly for LTR colonized with *Pseudomonas* or other pathogenic bacterial taxa. With the advancement in knowledge in host-microbial interactions in the lung, the ability to develop targeted treatments of dysregulated host-microbiome pathways could improve the effects of allograft dysfunction.

We recognize our studies have limitations. Our findings represent proof-of principle in isolating critical mechanistic pathways of innate immune responses and warrant further translation into human models of lung allograft injury. Our use of the THP-1 macrophages to investigate the relationship between bacteria and macrophage IL-1β production was validated in human AM. However, our experiments detailing IL-1β production following stimulation with LTR BAL microbiota and our treatment of macrophages with inflammasome inhibitors after *PsA* stimulation will be strengthened by validation in human AM in future studies. Additionally, stimulations with LTR lung microbiota should be repeated with differing relative abundances of *Pseudomonas* to represent the heterogeneity of microbiome composition seen in LTR. While it is possible that IL-1β could be produced outside of the canonical inflammasome pathway, future studies will determine the inflammasome sensors activated by *PsA* and *Pseudomonas-dominated* lung microbiota. It was beyond the scope of this manuscript to examine the association to CLAD, as AMs were selected from healthy LTR and then exogenously exposed to microbiota. Future studies that longitudinally characterize AM responses from a heterogeneous mix of LTR to their microbiota and to lung allograft dysfunction would further our understanding of this mechanistic pathway in the development of CLAD. Finally, the use of inflammasome inhibitors on other macrophage biology will need to be assessed, as the potential anti-inflammatory benefits would need to outweigh impairment in the ability to fight off life-threatening infections.

In conclusion, we observed a sustained increase in the production of IL-1β by human LTR -derived AM stimulated with *PsA* compared to other bacteria. Stimulation with *PsA* dominant lung microbiota from LTR induced upregulation of IL-1β in THP-1 macrophages. *PsA*-induced macrophage IL-1β production was abrogated with inflammasome inhibitor treatment. Our findings provide evidence for inflammasome activation as a potential mechanistic link between *Pseudomonas* and *Pseudomonas-dominated* lung microbiome dysbiosis and sustained innate immune activation by AM in otherwise healthy LTR.

## Acknowledgments

Microbial culturing was performed at the University of Pennsylvania’s Microbial Culture & Metabolomics Core which is supported by the Center for Molecular Studies in Digestive & Liver Diseases (P30DK050306). The PennCHOP Microbiome Program performed the 16S rRNA sequencing. This work was supported by NIH grants 5T32HL007534-38 (NB), R33-HL137062, and R01-HL113252 (RGC), U01 HL145435 (PS) and Cystic Fibrosis Foundation grant Christ18AB0.

