## Supplemental Data for "*Pseudomonas aeruginosa* Elicits Sustained IL-1β Upregulation in Alveolar Macrophages from Lung Transplant Recipients"

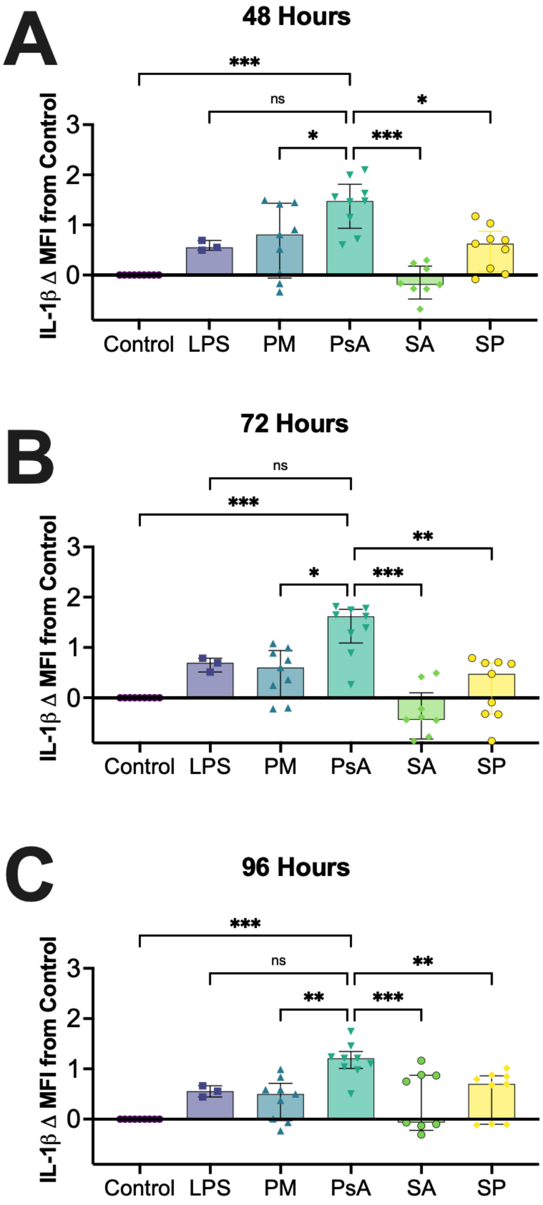


**Supplementary Figure 1:** *THP-1MΦ express elevated IL-1β in response to PsA compared to other bacteria.* THP-1MΦ (n=18) were stimulated with heat inactivated bacteria at a multiplicity of infection of 1:1 for 48 **(A)**, 72 **(B)** and 96 **(C)** hours. Cells were incubated with Golgi inhibitors the last 4 hours, stained for high dimensional flow cytometry. IL-1β is expressed as mean fluorescence intensity (MFI) fold change compared to expression in macrophages at baseline conditions (control). **ns** = not significant; * $p \leq 0.05$; ** $p \leq 0.002$; *** $p \leq 0.001$; **LPS**= lipopolysaccharide, **PM**= P. melaninogenica, **PsA**=P. aeruginosa, **SA**= S. aureus, **SP**= S. pneumoniae

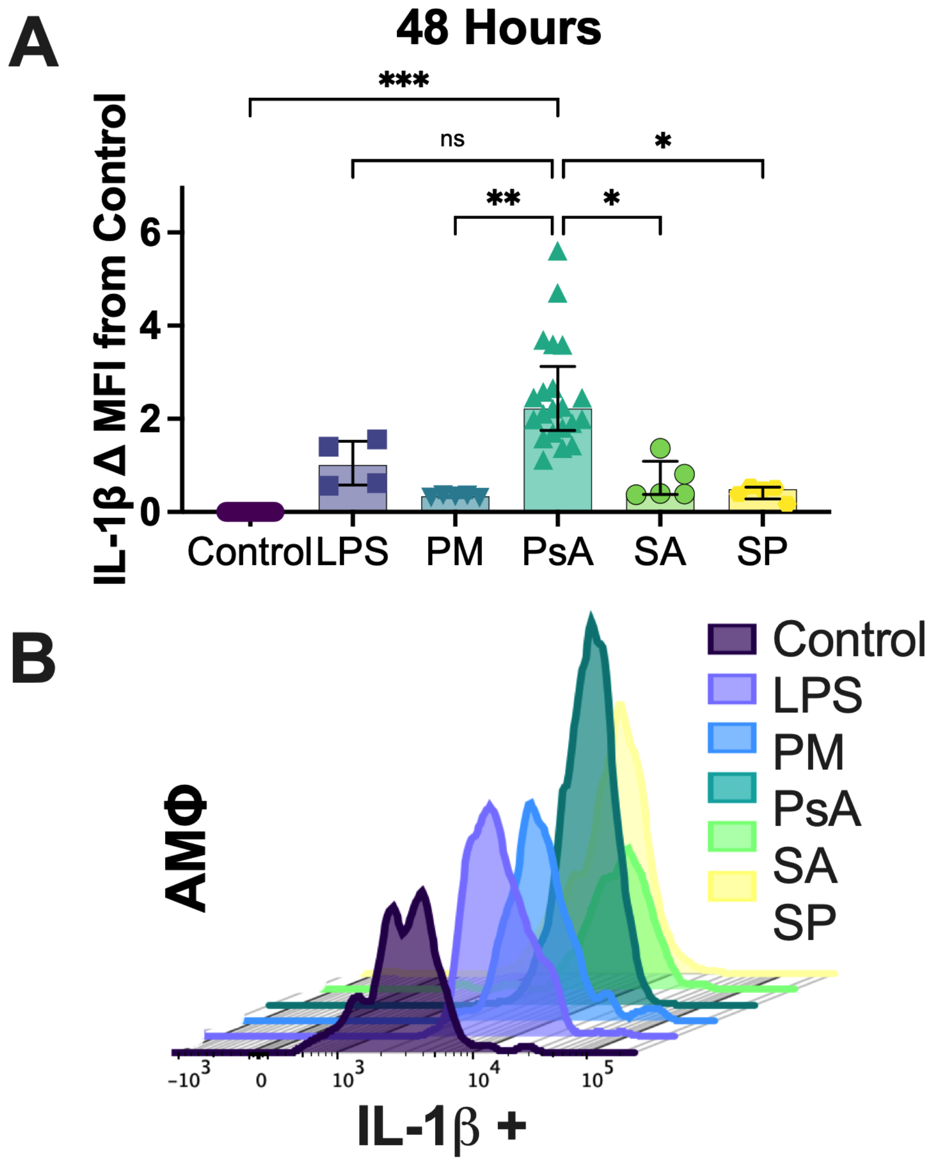


**Supplementary Figure 2 :** *AMΦ responses to different bacteria at 48 hours. AMΦ* were stimulated with heat inactivated bacteria at a multiplicity of infection of 1:1 for 48 hours. Cells were incubated with Golgi inhibitors the last 4 hours, stained for high dimensional flow cytometry IL-1β is expressed as mean fluorescence intensity (MFI) fold change compared to expression in macrophages at baseline conditions (control). **A)** *AMΦ* express elevated IL-1β in response to *PsA* compared to other bacteria. **B)** Representative histograms demonstrating IL-1β expression after bacterial stimulation for 48 hours. **LPS** (n=5), **PM** (n=5), **PsA** (n=20), **SA** (n=5), **SP** (n=5); **ns** = not significant; * p ≤0.05; ** p ≤0.002; *** p ≤ 0.001; **LPS**= lipopolysaccharide, **PM**= P. melaninogenica, **PsA**=P. aeruginosa, **SA**= S. aureus, **SP**= S. pneumoniae

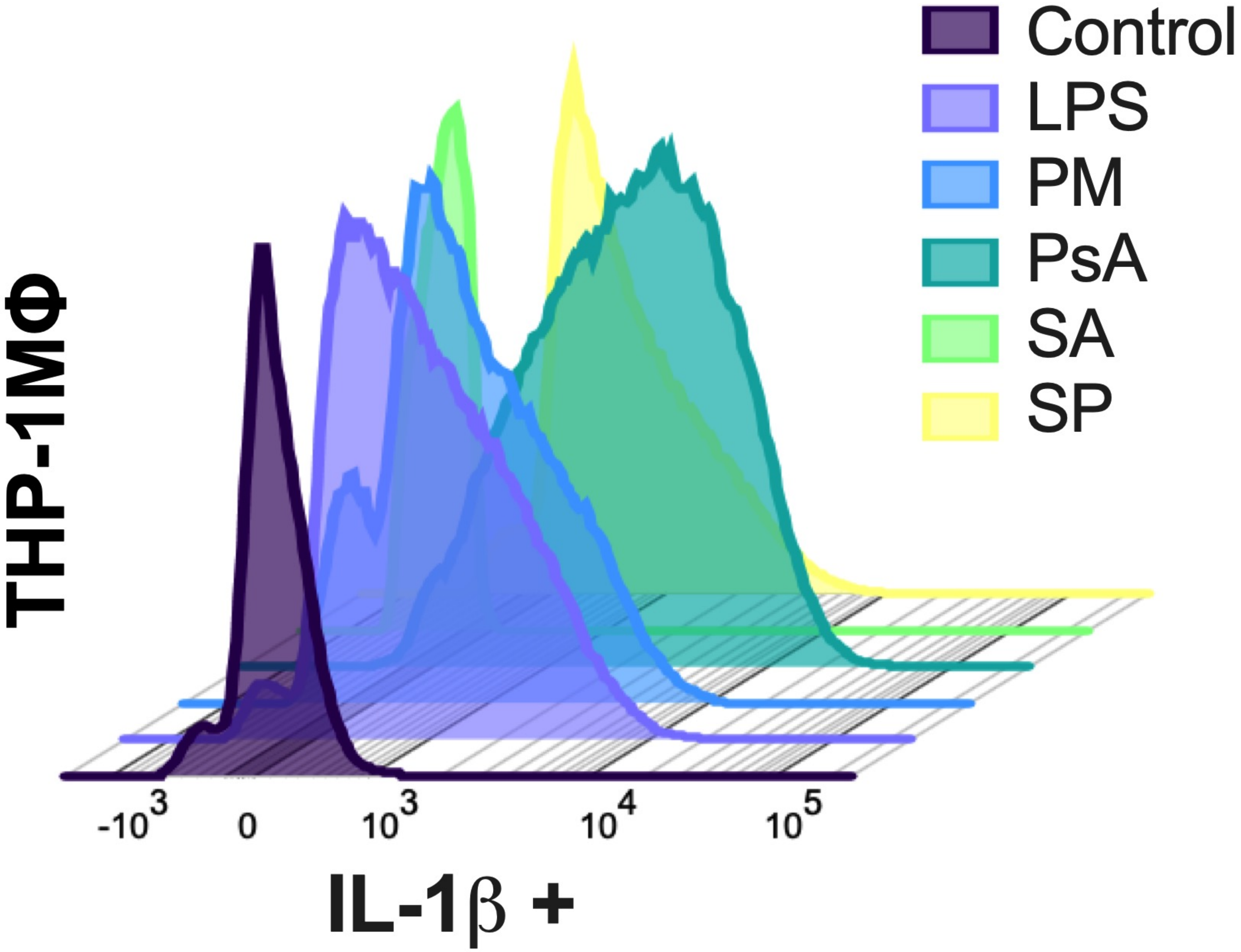


**Supplementary Figure 3:** *THP-1MΦ responses to different bacteria at 24 hours.* THP-1MΦ (n=18) were stimulated with heat inactivated bacteria at a multiplicity of infection of 1:1 for 24hours. Cells were incubated with Golgi inhibitors the last 4 hours, stained for high dimensional flow cytometry. Representative histogram of IL-1β fluorescent intensity from flow cytometry. **LPS**= lipopolysaccharide, **PM**= P. melaninogenica, **PsA**=P. aeruginosa, **SA**= S. aureus, **SP**= S. pneumoniae

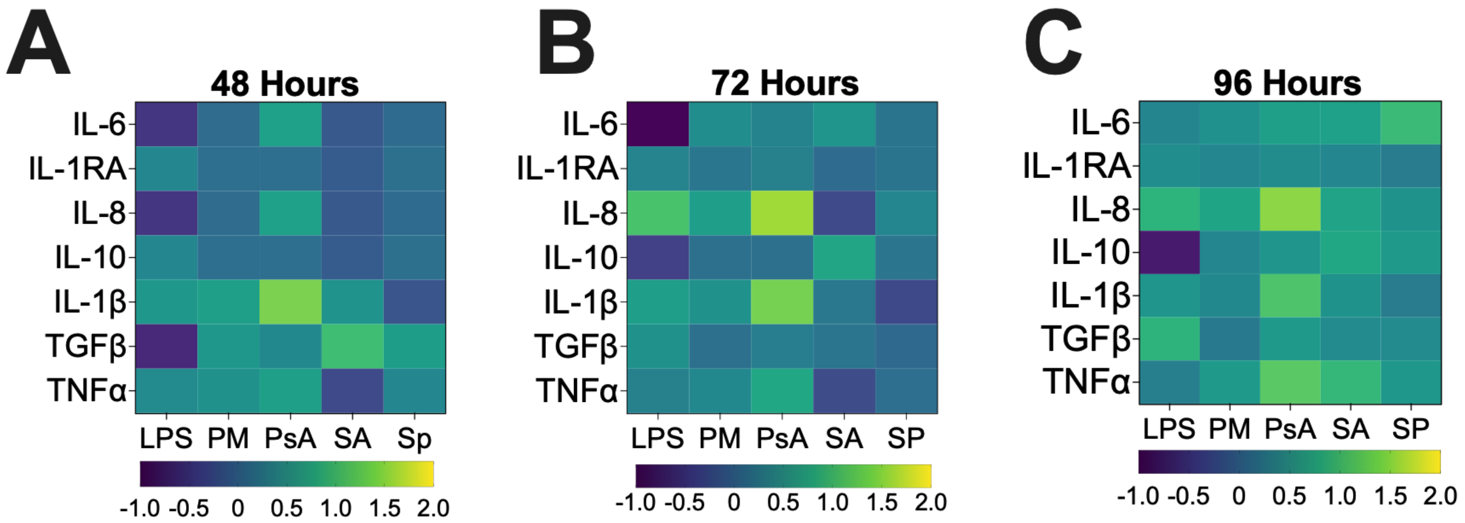


**Supplementary Figure 4:** *THP-1MΦ responses to different bacteria over time.* THP-1MΦ (n=18) were stimulated with heat inactivated bacteria at a multiplicity of infection of 1:1 for 48 **(A)**, 72 **(B)** and 96 **(C)** hours. Cells were incubated with Golgi inhibitors the last 4 hours, stained for high dimensional flow cytometry. Each marker is expressed as mean fluorescence intensity (MFI) fold change compared to expression in macrophages at baseline conditions (control).

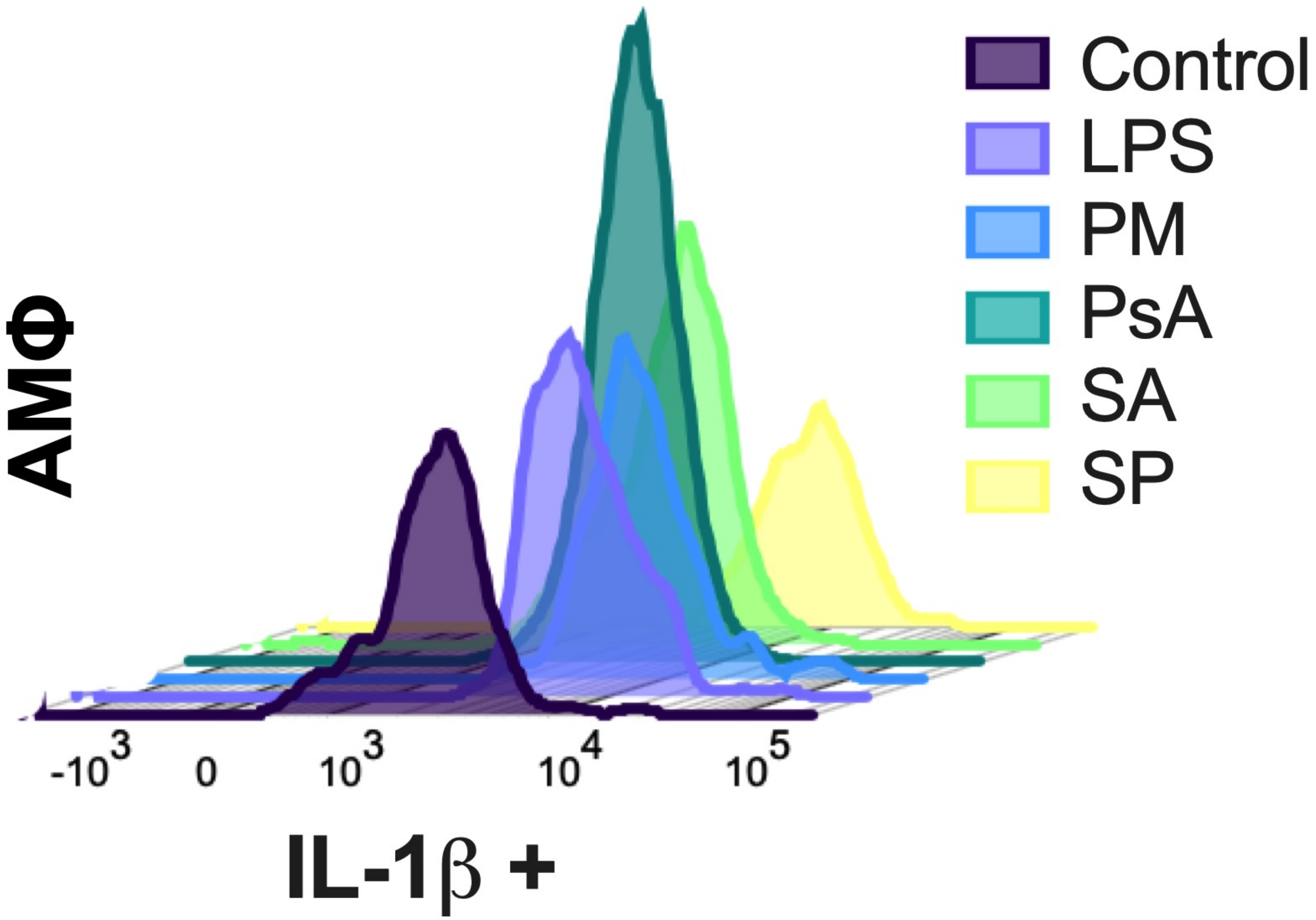


**Supplementary Figure :** *AMΦ responses to different bacteria at 24 hours. AMΦ* were stimulated with heat inactivated bacteria at a multiplicity of infection of 1:1 for 24 hours. Cells were incubated with Golgi inhibitors the last 4 hours, stained for high dimensional flow cytometry. Representative histograms demonstrating IL-1β expression after bacterial stimulation for 48 hours. **LPS** (n=17), **PM** (n=10), **PsA** (n=20), **SA** (n=5), **SP** (n=10); **LPS**= lipopolysaccharide, **PM**= P. melaninogenica, **PsA**=P. aeruginosa, **SA**= S. aureus, **SP**= S. pneumoniae

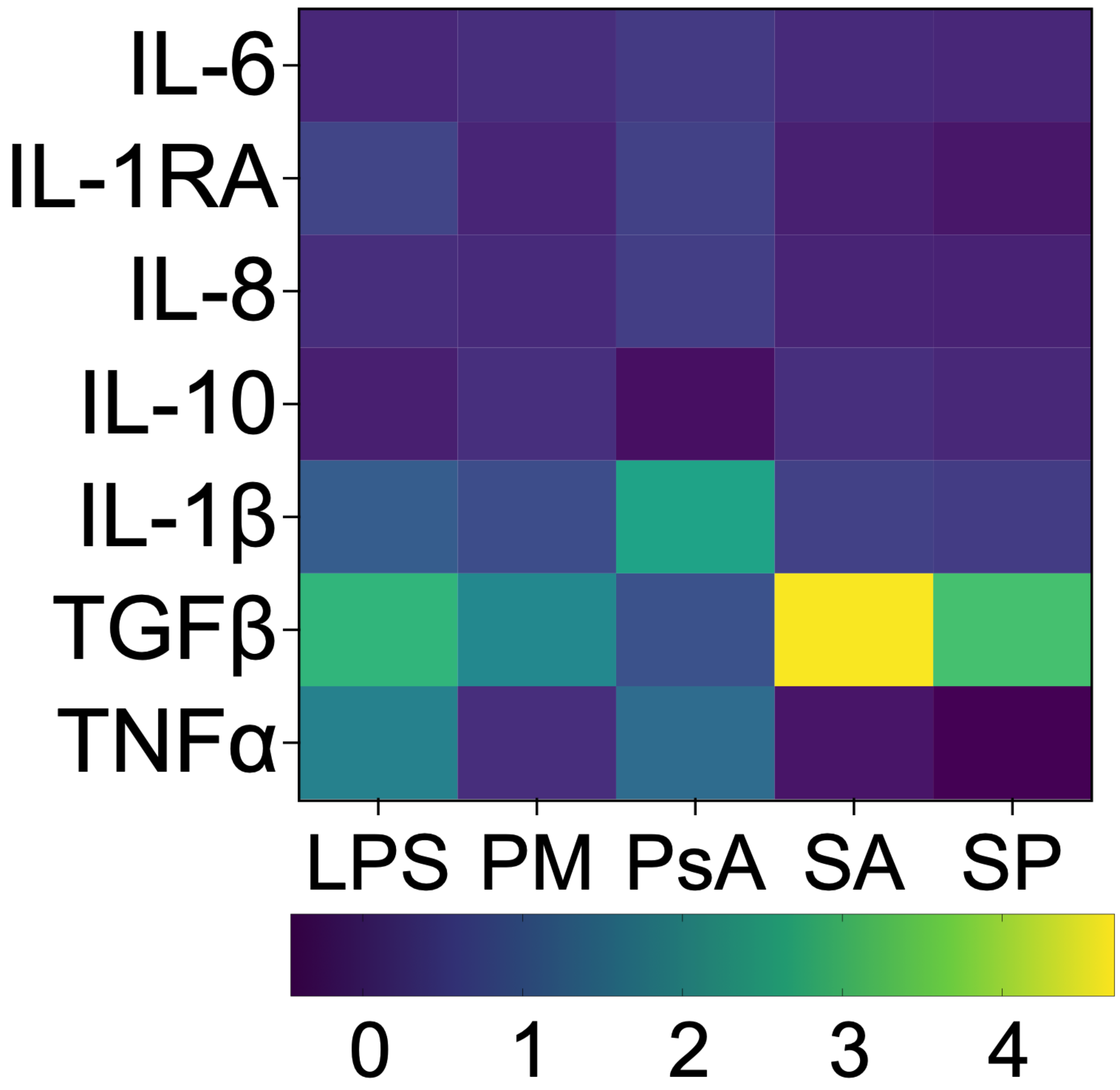


**Supplementary Figure XXX:** *AMΦ responses to different bacteria at 48 hours.* AMΦ were stimulated with heat inactivated bacteria at a multiplicity of infection of 1:1 for 48 hours. Cells were incubated with Golgi inhibitors the last 4 hours, stained for high dimensional flow cytometry. Each marker is expressed as mean fluorescence intensity (MFI) fold change compared to expression in macrophages at baseline conditions (control). **LPS** (n=5), **PM** (n=5), **PsA** (n=20), **SA** (n=5), **SP** (n=5); **LPS**= lipopolysaccharide, **PM**= P. melaninogenica, **PsA**=P. aeruginosa, **SA**= S. aureus, **SP**= S. pneumoniae

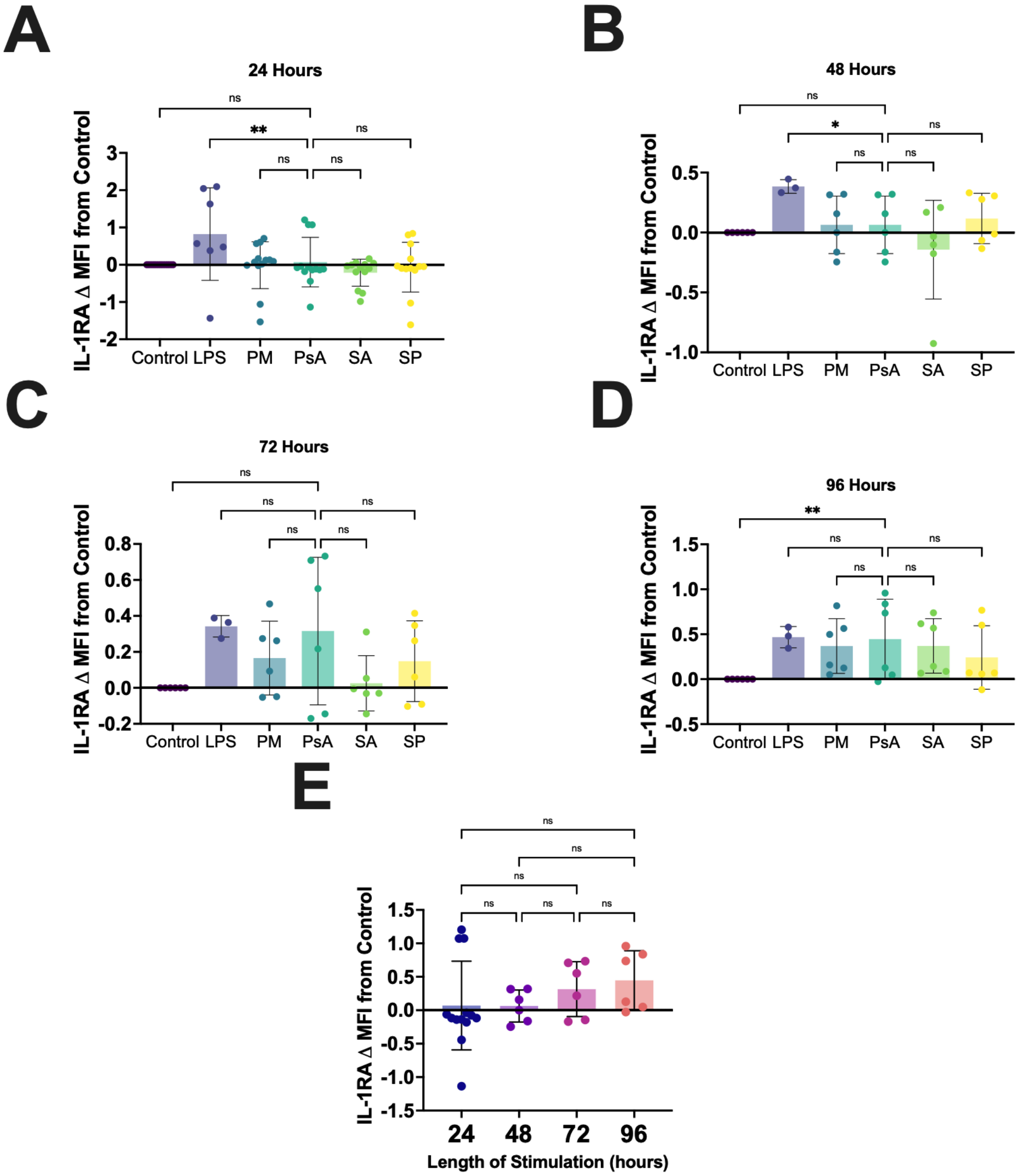


**Supplementary Figure 5:** *THP-1MΦ IL-1RA expression in response to different bacteria.* THP-1MΦ (n=18) were stimulated with heat inactivated bacteria at a multiplicity of infection of 1:1 for 24 **(A)**, 48 **(B)**, 72 **(C)** and 96 **(D)** hours. Cells were incubated with Golgi inhibitors the last 4 hours, stained for high dimensional flow cytometry. IL-1RA is expressed as mean fluorescence intensity (MFI) fold change compared to expression in macrophages at baseline conditions (control). **E)** IL-1RA kinetics after challenge with *PsA* for up to 96 hours. **ns** = not significant; * p ≤0.05; ** p ≤0.002; *** p ≤ 0.001; **LPS**= lipopolysaccharide, **PM**= P. melaninogenica, **PsA**=P. aeruginosa, **SA**= S. aureus, **SP**= S. pneumoniae

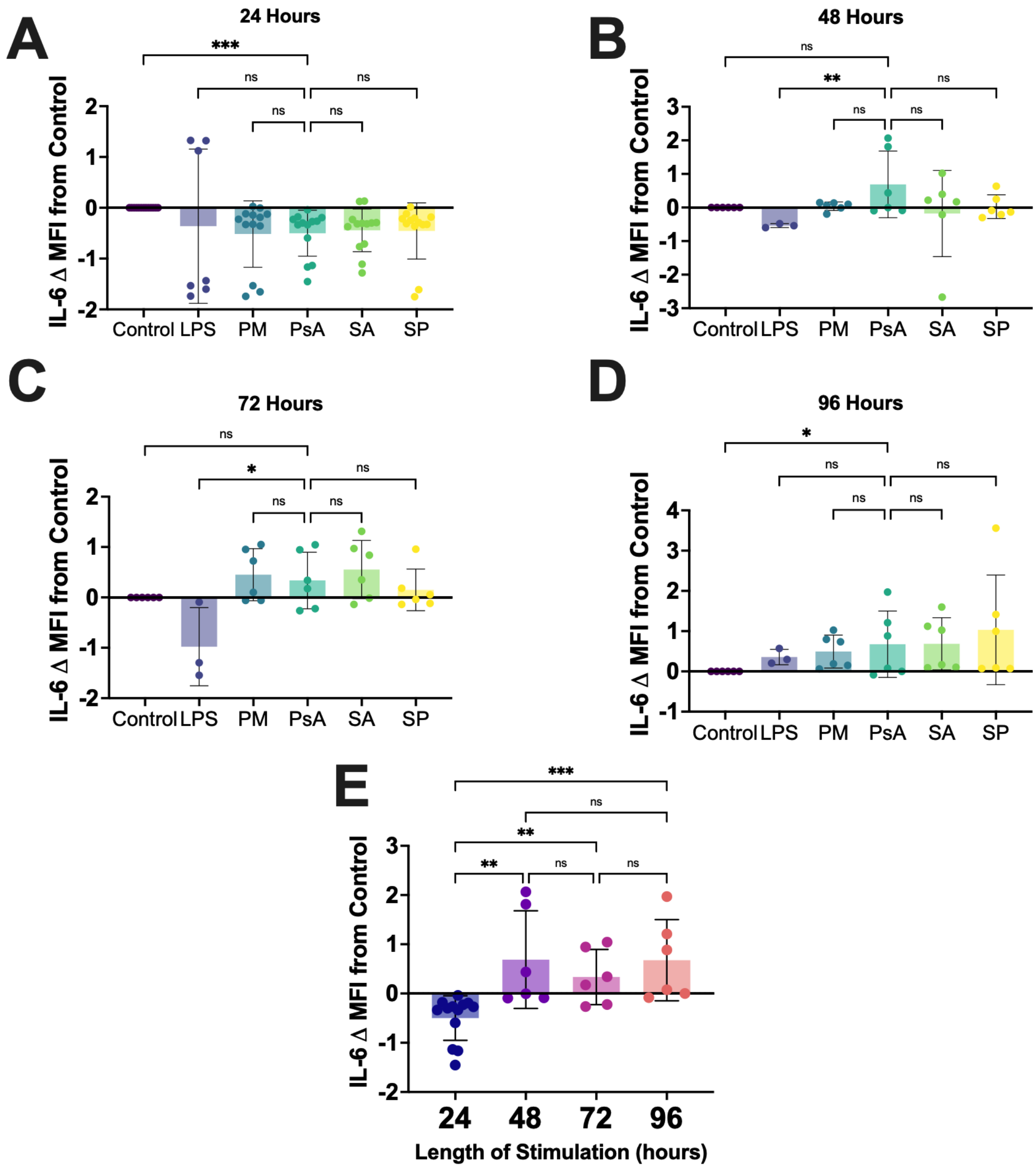


**Supplementary Figure 6:** *THP-1MΦ IL-6 expression in response to different bacteria.* THP-1MΦ (n=18) were stimulated with heat inactivated bacteria at a multiplicity of infection of 1:1 for 24 **(A)**, 48 **(B)**, 72 **(C)** and 96 **(D)** hours. Cells were incubated with Golgi inhibitors the last 4 hours, stained for high dimensional flow cytometry. IL-6 is expressed as mean fluorescence intensity (MFI) fold change compared to expression in macrophages at baseline conditions (control). **E)** IL-6 kinetics after challenge with *PsA* for up to 96 hours. **ns** = not significant; * $p \leq 0.05$; ** $p \leq 0.002$; *** $p \leq 0.001$; **LPS**= lipopolysaccharide, **PM**= P. melaninogenica, **PsA**=P. aeruginosa, **SA**= S. aureus, **SP**= S. pneumoniae

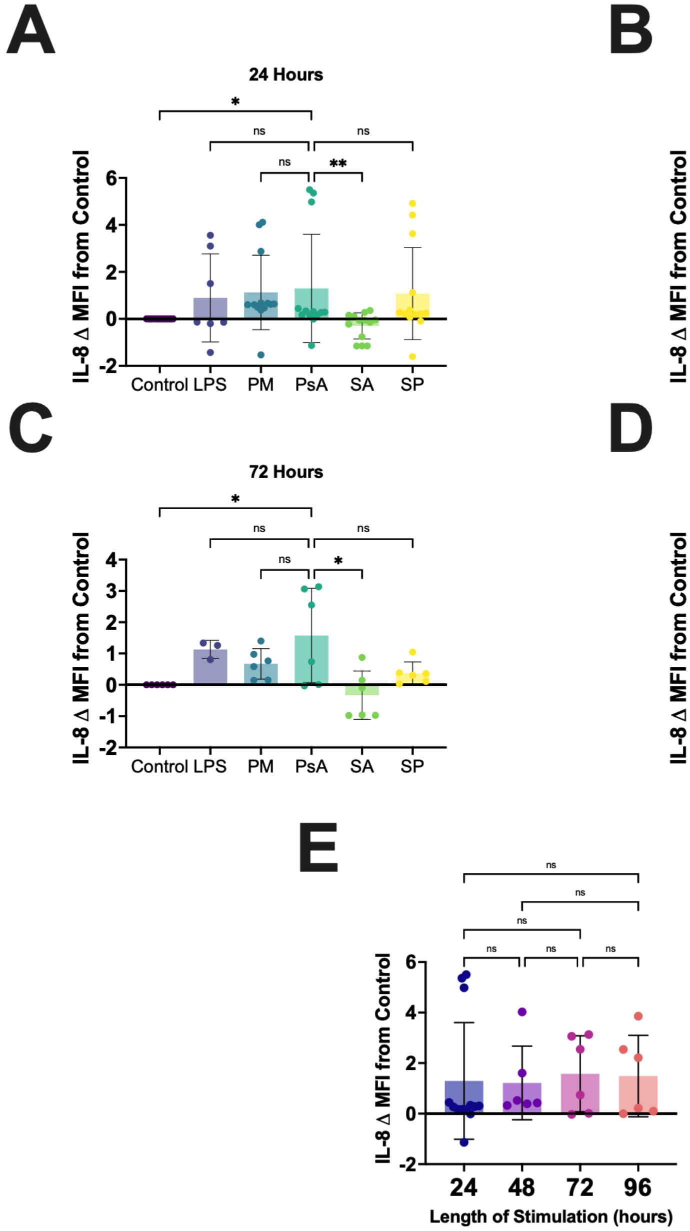


**Supplementary Figure 7:** *THP-1MΦ IL-8 expression in response to different bacteria.* THP-1MΦ (n=18) were stimulated with heat inactivated bacteria at a multiplicity of infection of 1:1 for 24 **(A)**, 48 **(B)**, 72 **(C)** and 96 **(D)** hours. Cells were incubated with Golgi inhibitors the last 4 hours, stained for high dimensional flow cytometry. IL-8 is expressed as mean fluorescence intensity (MFI) fold change compared to expression in macrophages at baseline conditions (control). **E)** IL-8 kinetics after challenge with *PsA* for up to 96 hours. **ns** = not significant; * $p \leq 0.05$; ** $p \leq 0.002$; *** $p \leq 0.001$; **LPS**= lipopolysaccharide, **PM**= P. melaninogenica, **PsA**=P. aeruginosa, **SA**= S. aureus, **SP**= S. pneumoniae

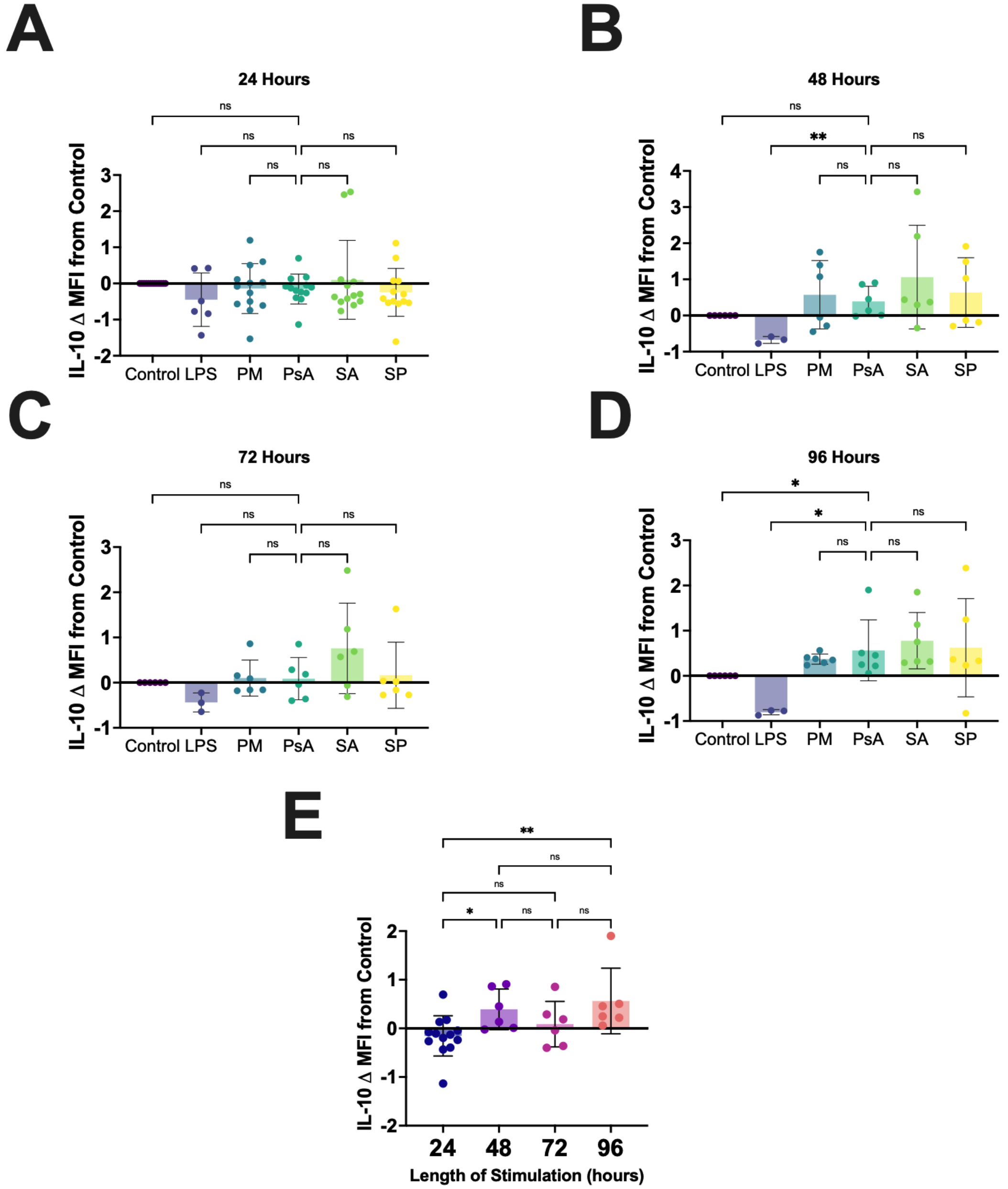


**Supplementary Figure 8:** *THP-1MΦ IL-10 expression in response to different bacteria.* THP-1MΦ (n=18) were stimulated with heat inactivated bacteria at a multiplicity of infection of 1:1 for 24 **(A)**, 48 **(B)**, 72 **(C)** and 96 **(D)** hours. Cells were incubated with Golgi inhibitors the last 4 hours, stained for high dimensional flow cytometry. IL-10 is expressed as mean fluorescence intensity (MFI) fold change compared to expression in macrophages at baseline conditions (control). **E)** IL-10 kinetics after challenge with *PsA* for up to 96 hours. **ns** = not significant; * $p \leq 0.05$; ** $p \leq 0.002$; *** $p \leq 0.001$; **LPS**= lipopolysaccharide, **PM**= P. melaninogenica, **PsA**=P. aeruginosa, **SA**= S. aureus, **SP**= S. pneumoniae

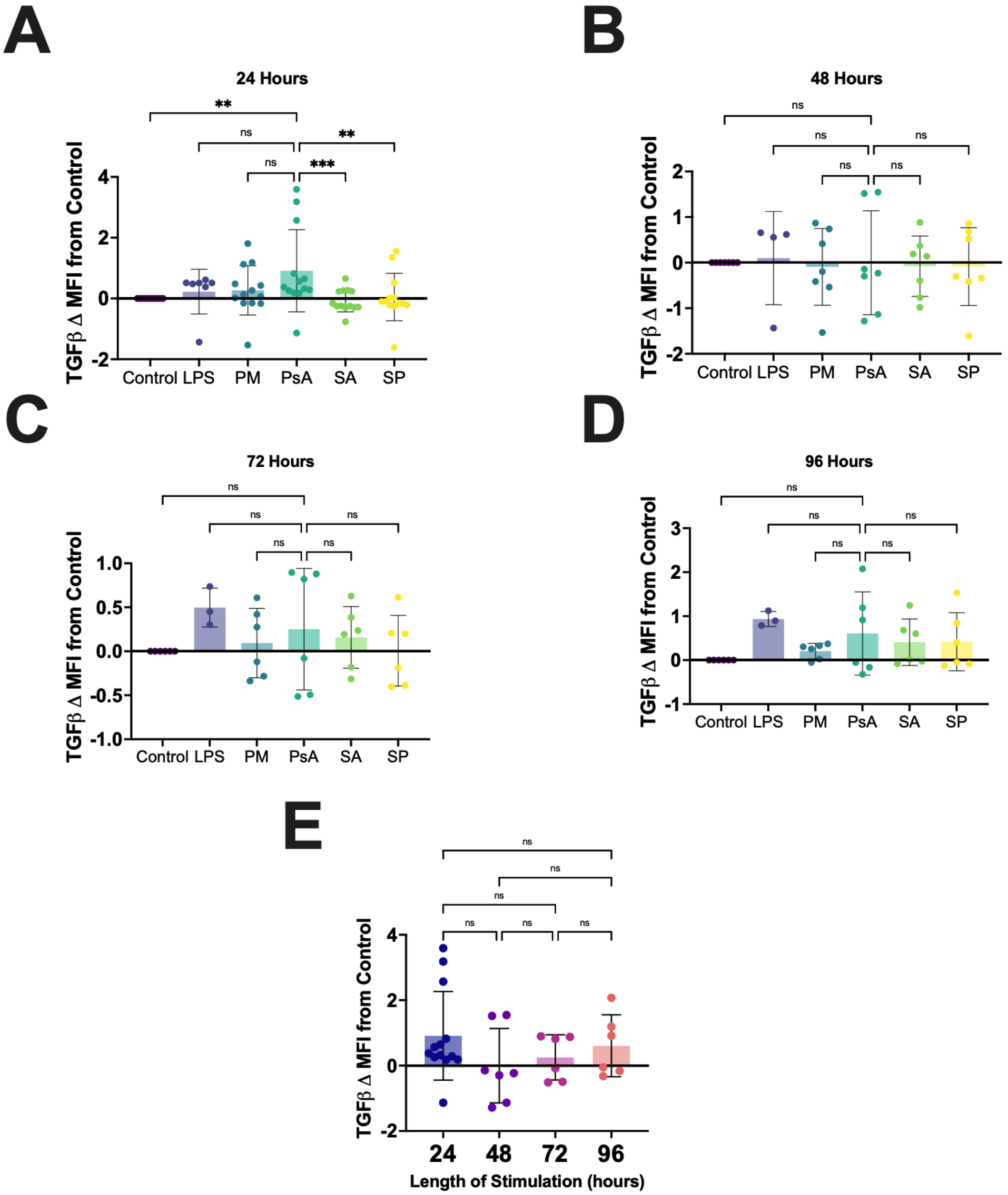


**Supplementary Figure 9:** *THP-1MΦ TGFβ expression in response to different bacteria.* THP-1MΦ (n=18) were stimulated with heat inactivated bacteria at a multiplicity of infection of 1:1 for 24 **(A)**, 48 **(B)**, 72 **(C)** and 96 **(D)** hours. Cells were incubated with Golgi inhibitors the last 4 hours, stained for high dimensional flow cytometry. TGFβ is expressed as mean fluorescence intensity (MFI) fold change compared to expression in macrophages at baseline conditions (control). **E)** TGFβ kinetics after challenge with *PsA* for up to 96 hours. **ns** = not significant; * p ≤0.05; ** p ≤0.002; *** p ≤ 0.001; **LPS**= lipopolysaccharide, **PM**= P. melaninogenica, **PsA**=P. aeruginosa, **SA**= S. aureus, **SP**= S. pneumoniae

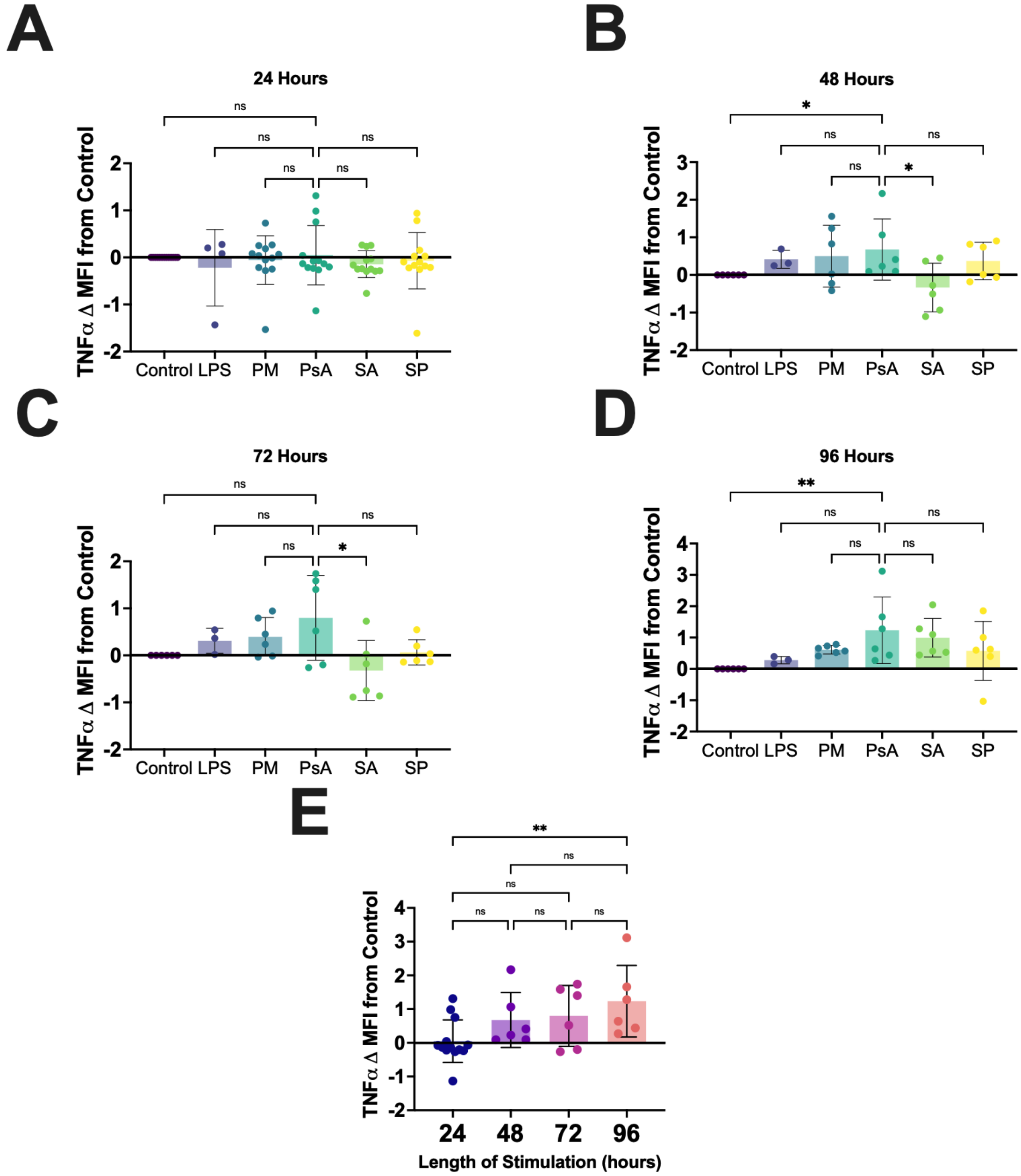


**Supplementary Figure 10:** *THP-1MΦ TNFα expression in response to different bacteria.* THP-1MΦ (n=18) were stimulated with heat inactivated bacteria at a multiplicity of infection of 1:1 for 24 **(A)**, 48 **(B)**, 72 **(C)** and 96 **(D)** hours. Cells were incubated with Golgi inhibitors the last 4 hours, stained for high dimensional flow cytometry. TNFα is expressed as mean fluorescence intensity (MFI) fold change compared to expression in macrophages at baseline conditions (control). **E)** TNFα kinetics after challenge with *PsA* for up to 96 hours. **ns** = not significant; * $p \leq 0.05$; ** $p \leq 0.002$; *** $p \leq 0.001$; **LPS**= lipopolysaccharide, **PM**= P. melaninogenica, **PsA**=P. aeruginosa, **SA**= S. aureus, **SP**= S. pneumoniae

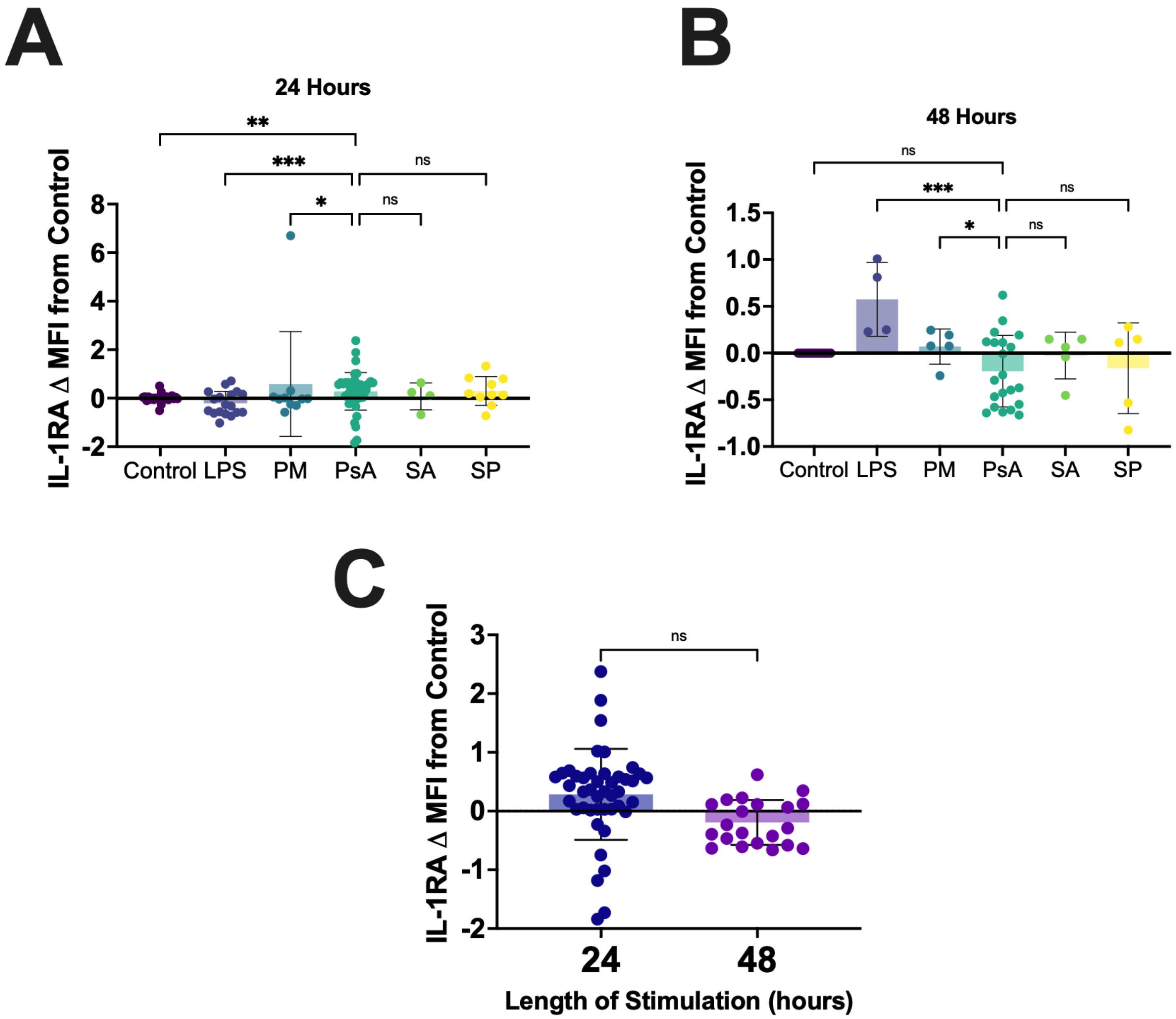


**Supplementary Figure 11:** *AMΦ IL-1RA expression in response to different bacteria.* AMΦ were stimulated with heat inactivated bacteria at a multiplicity of infection of 1:1 for 24 **(A)** and 48 **(B)** hours. Cells were incubated with Golgi inhibitors the last 4 hours, stained for high dimensional flow cytometry. IL-1RA is expressed as mean fluorescence intensity (MFI) fold change compared to expression in macrophages at baseline conditions (control). **C)** IL-1RA kinetics after challenge with *PsA* for up to 48 hours. **LPS** (n=5), **PM** (n=5), **PsA** (n=20), **SA** (n=5), **SP** (n=5); **ns** = not significant; * $p \leq 0.05$; ** $p \leq 0.002$; *** $p \leq 0.001$; **LPS**= lipopolysaccharide, **PM**= P. melaninogenica, **PsA**=P. aeruginosa, **SA**= S. aureus, **SP**= S. pneumoniae

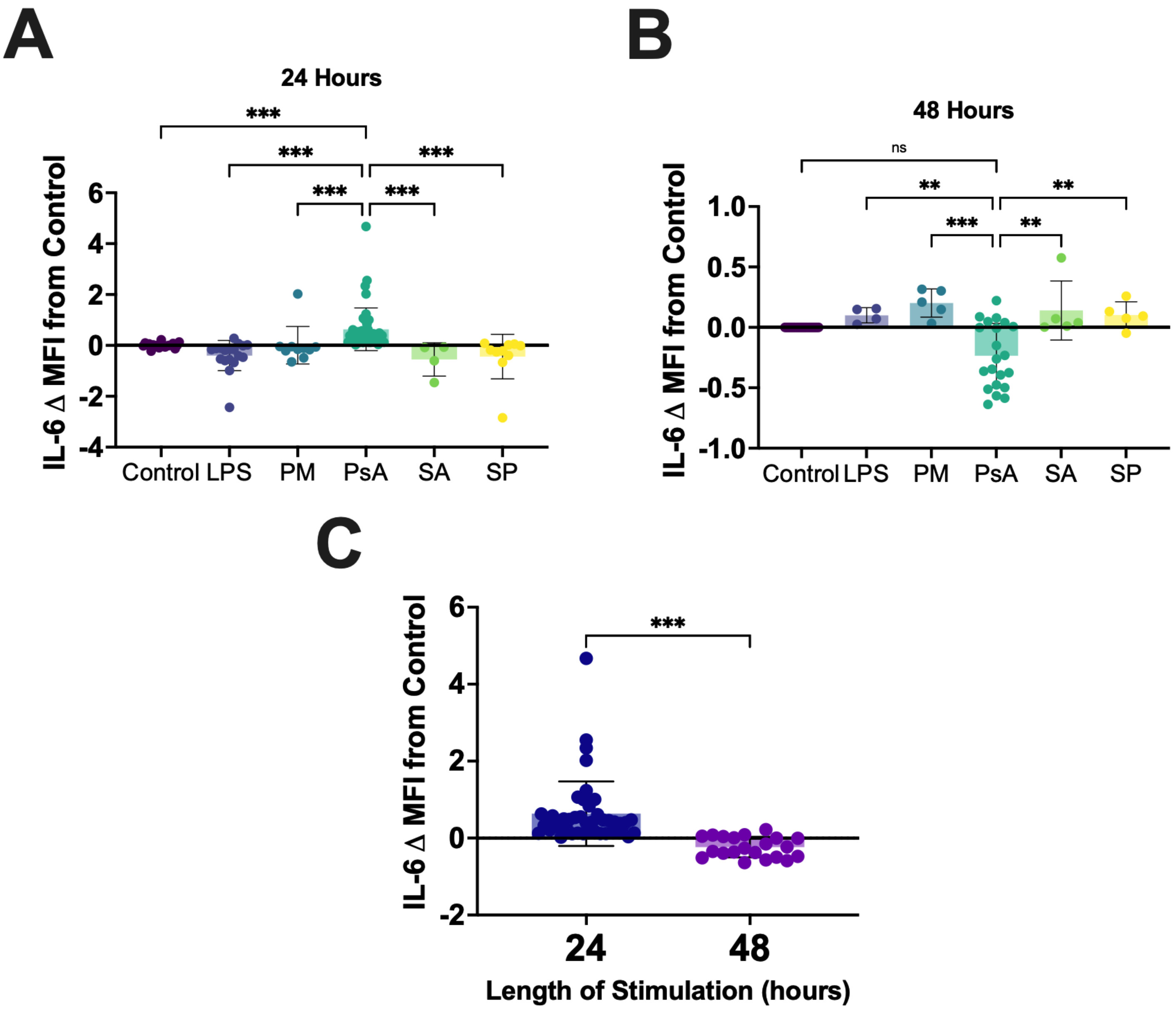


**Supplementary Figure 12:** *AMΦ IL-6 expression in response to different bacteria.* AMΦ were stimulated with heat inactivated bacteria at a multiplicity of infection of 1:1 for 24 **(A)** and 48 **(B)** hours. Cells were incubated with Golgi inhibitors the last 4 hours, stained for high dimensional flow cytometry. IL-6 is expressed as mean fluorescence intensity (MFI) fold change compared to expression in macrophages at baseline conditions (control). **C)** IL-6 kinetics after challenge with *PsA* for up to 48 hours. **LPS** (n=5), **PM** (n=5), **PsA** (n=20), **SA** (n=5), **SP** (n=5); **ns** = not significant; * $p \leq 0.05$; ** $p \leq 0.002$; *** $p \leq 0.001$; **LPS**= lipopolysaccharide, **PM**= P. melaninogenica, **PsA**=P. aeruginosa, **SA**= S. aureus, **SP**= S. pneumoniae

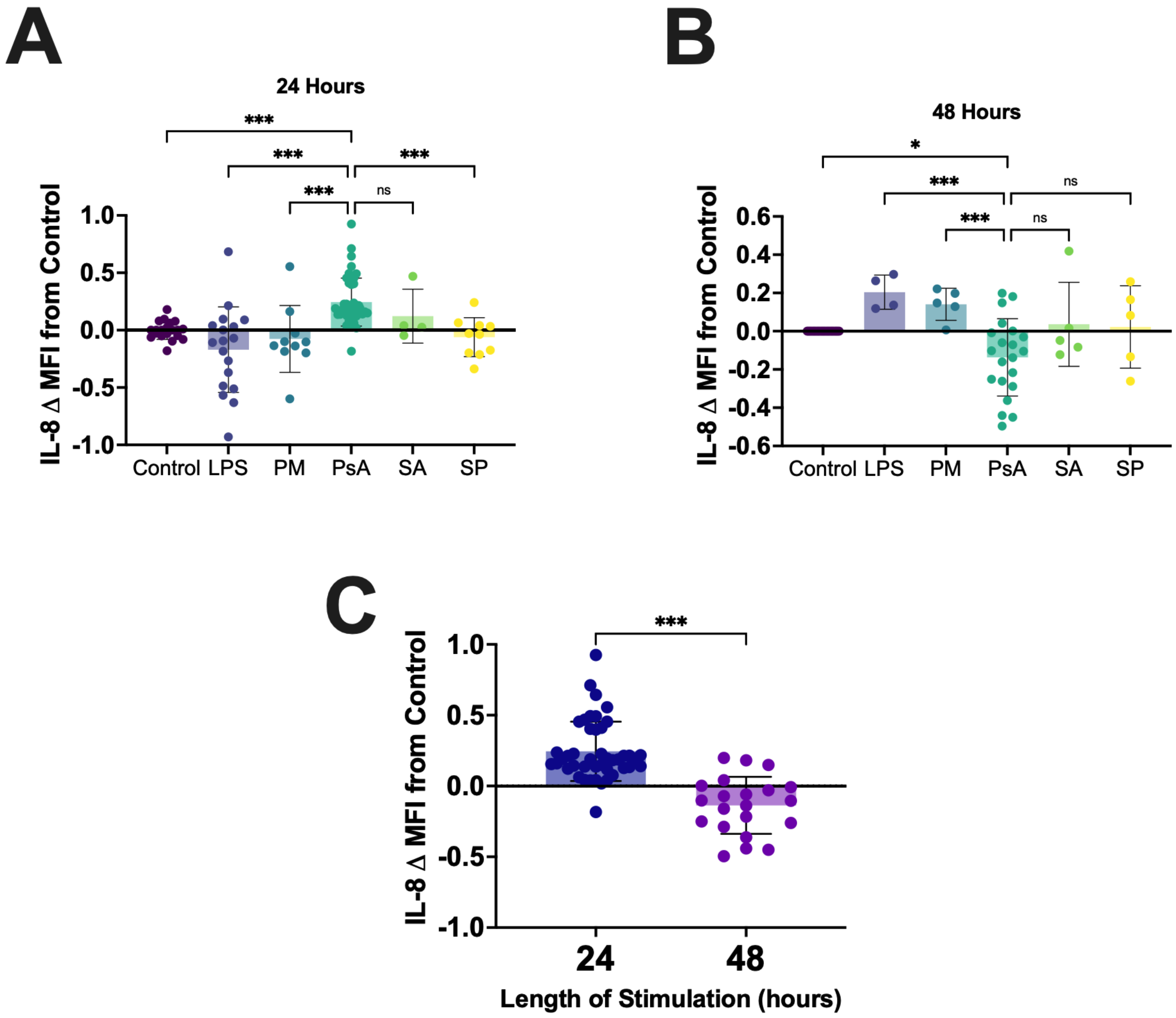


**Supplementary Figure 13:** *AMΦ IL-8 expression in response to different bacteria.* AMΦ were stimulated with heat inactivated bacteria at a multiplicity of infection of 1:1 for 24 **(A)** and 48 **(B)** hours. Cells were incubated with Golgi inhibitors the last 4 hours, stained for high dimensional flow cytometry. IL-8 is expressed as mean fluorescence intensity (MFI) fold change compared to expression in macrophages at baseline conditions (control). **C)** IL-8 kinetics after challenge with *PsA* for up to 48 hours. **LPS** (n=5), **PM** (n=5), **PsA** (n=20), **SA** (n=5), **SP** (n=5); **ns** = not significant; * p ≤0.05; ** p ≤0.002; *** p ≤ 0.001; **LPS**= lipopolysaccharide, **PM**= P. melaninogenica, **PsA**=P. aeruginosa, **SA**= S. aureus, **SP**= S. pneumoniae

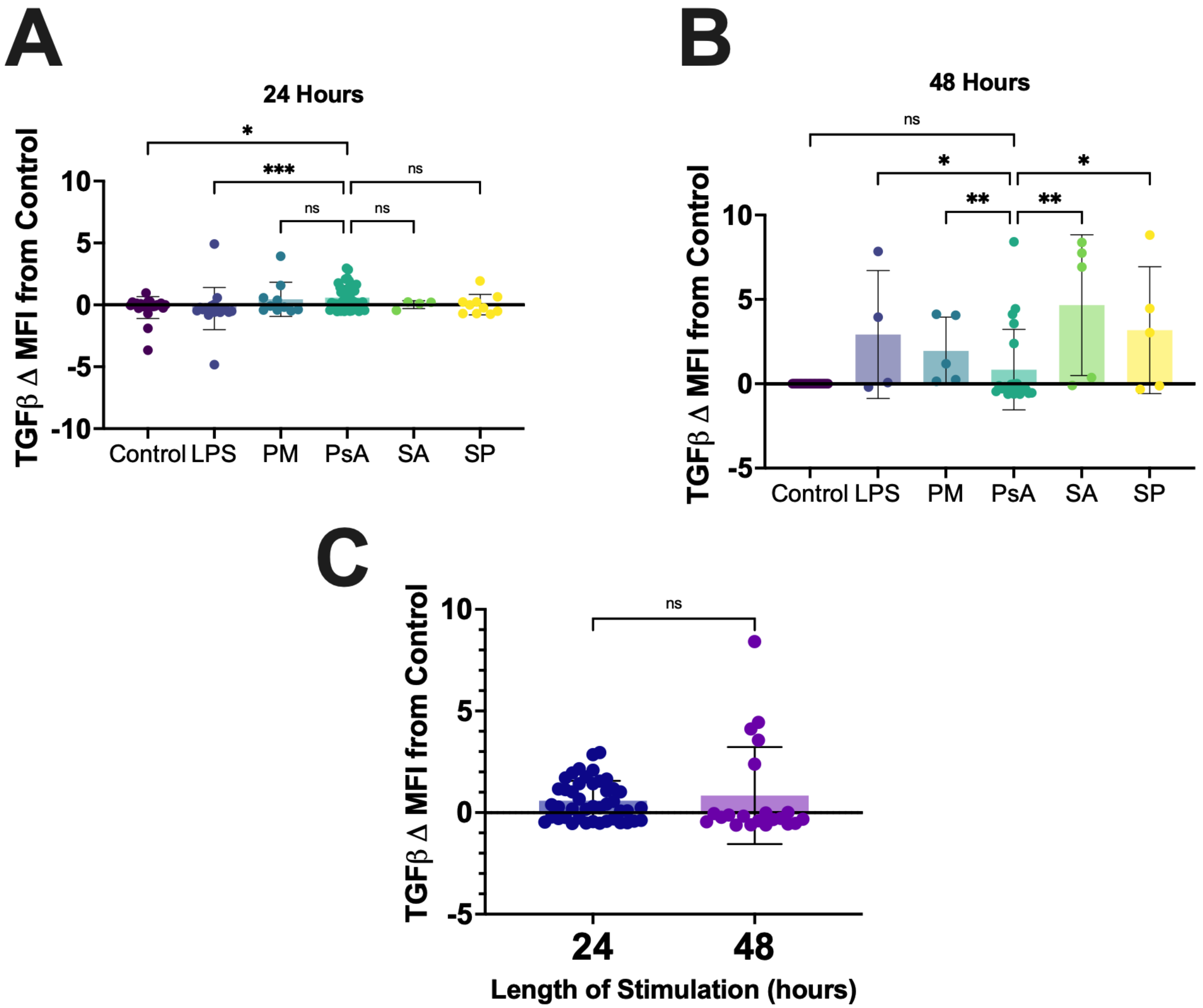


**Supplementary Figure 14:** *AMΦ TGFβ expression in response to different bacteria.* AMΦ were stimulated with heat inactivated bacteria at a multiplicity of infection of 1:1 for 24 **(A)** and 48 **(B)** hours. Cells were incubated with Golgi inhibitors the last 4 hours, stained for high dimensional flow cytometry. TGFβ is expressed as mean fluorescence intensity (MFI) fold change compared to expression in macrophages at baseline conditions (control). **C)** TGFβ kinetics after challenge with *PsA* for up to 48 hours. **LPS** (n=5), **PM** (n=5), **PsA** (n=20), **SA** (n=5), **SP** (n=5); **ns** = not significant; * p ≤0.05; ** p ≤0.002; *** p ≤ 0.001; **LPS**= lipopolysaccharide, **PM**= P. melaninogenica, **PsA**=P. aeruginosa, **SA**= S. aureus, **SP**= S. pneumoniae

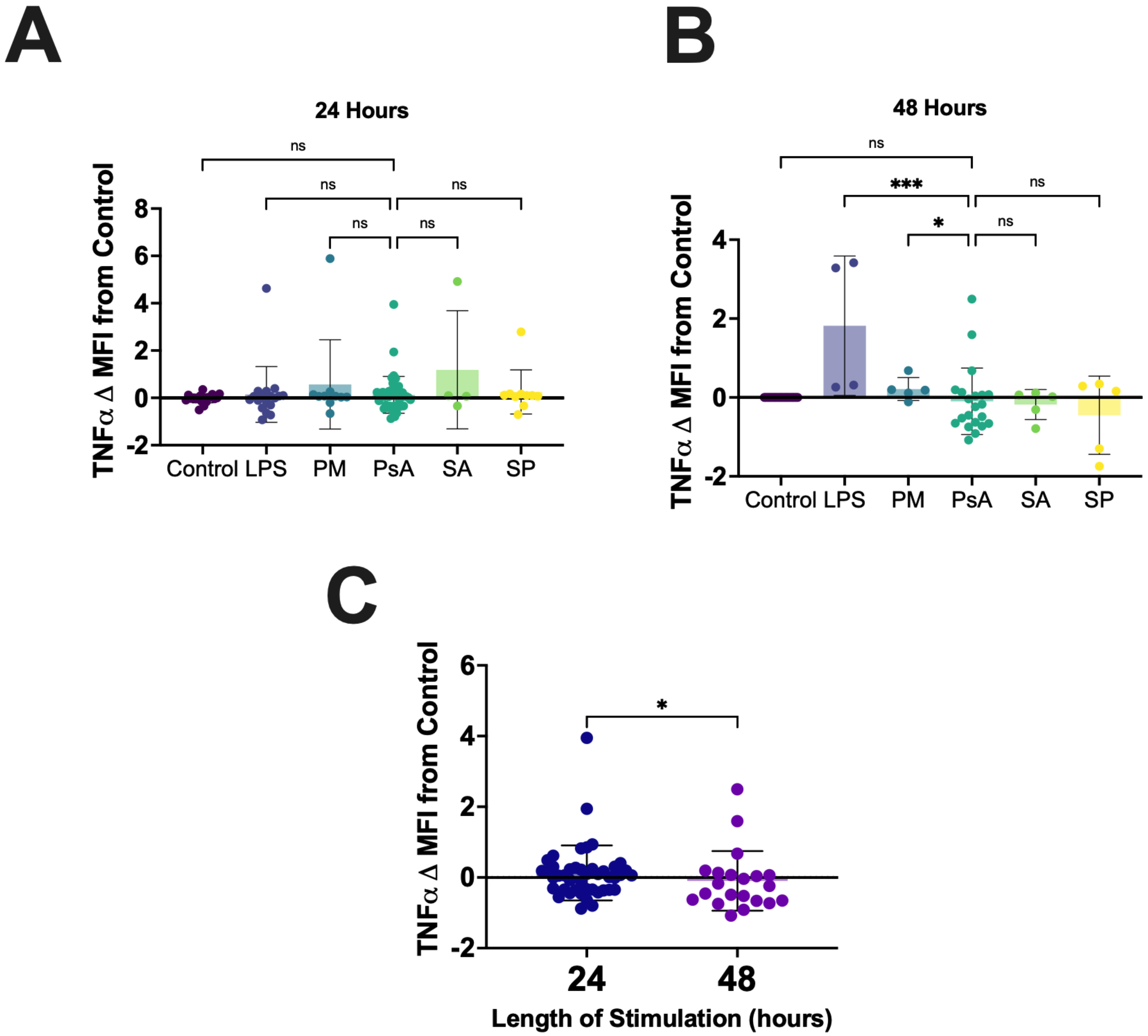


**Supplementary Figure 15:** *AMΦ TNFα expression in response to different bacteria.* AMΦ were stimulated with heat inactivated bacteria at a multiplicity of infection of 1:1 for 24 **(A)** and 48 **(B)** hours. Cells were incubated with Golgi inhibitors the last 4 hours, stained for high dimensional flow cytometry. TNFα is expressed as mean fluorescence intensity (MFI) fold change compared to expression in macrophages at baseline conditions (control). **C)** TNFα kinetics after challenge with *PsA* for up to 48 hours. **LPS** (n=5), **PM** (n=5), **PsA** (n=20), **SA** (n=5), **SP** (n=5); **ns** = not significant; * $p \leq 0.05$; ** $p \leq 0.002$; *** $p \leq 0.001$; **LPS**= lipopolysaccharide, **PM**= P. melaninogenica, **PsA**=P. aeruginosa, **SA**= S. aureus, **SP**= S. pneumoniae
